# Diversity and Function of Motile Ciliated Cell Types within Ependymal Lineages of the Zebrafish Brain

**DOI:** 10.1101/2021.02.17.431442

**Authors:** Percival P. D’Gama, Tao Qiu, Mehmet Ilyas Cosacak, Yan Ling Chong, Ahsen Konac, Jan Niklas Hansen, Christa Ringers, Subhra P. Hui, Emilie W. Olstad, Chee Peng Ng, Dheeraj Rayamajhi, Dagmar Wachten, David Liebl, Kazu Kikuchi, Caghan Kizil, Emre Yaksi, Sudipto Roy, Nathalie Jurisch-Yaksi

## Abstract

Motile cilia defects impair cerebrospinal fluid (CSF) flow, and can cause brain and spine disorders. To date, the development of ciliated cells, their impact on CSF flow and their function in brain and axial morphogenesis are not fully understood. Here, we have characterized motile ciliated cells within the zebrafish brain ventricles. We show that the ventricular surface undergoes significant restructuring through development, involving a transition from mono- to multiciliated cells (MCCs) driven by gmnc. MCCs are translationally polarized, co-exist with monociliated cells and generate directional flow patterns. Moreover, these ciliated cells have different developmental origins, and are genetically heterogenous with respect to expression of the Foxj1 family of ciliary master regulators. Finally, we show that cilia loss from specific brain regions or global perturbation of multiciliation does not affect overall brain or spine morphogenesis, but results in enlarged ventricles. Our findings establish that motile ciliated cells are generated by complementary and sequential transcriptional programs to support ventricular development.

## INTRODUCTION

The ependyma, which is the cellular layer lining the surface of the brain ventricles and spinal canal, plays a key role in CSF dynamics [1]. In the adult mammalian brain, the ependyma is composed primarily of post-mitotic glia-like cells known as multiciliated ependymal cells [2–5]. Ependymal cells are derived from radial glial cells and differentiate during the perinatal and postnatal periods [4, 6, 7]. Like MCCs in other tissues [5, 8], ependymal MCCs carry bundles of motile cilia on their apical surface, which beat and contribute to directional CSF flow [9–12]. In analogy to mammals, the zebrafish ependyma is also decorated with motile ciliated cells [12–20]. These motile ciliated cells appear as early as 28-32 hours post fertilization (hpf) in the embryonic brain [13–15], and even earlier in the central canal of the spinal cord [21–23]. During these embryonic stages, the cells bear solitary cilia that move in a rotational manner and generate a directional flow [14, 23]. Ependymal MCCs, analogous to mammalian ependymal cells, have only been reported in the adult zebrafish brain [16–20]. Yet, it remains unclear when and where these MCCs are established and how they contribute to CSF dynamics.

Besides their role in CSF circulation, ependymal MCCs are also necessary for the adult neurogenic niche to assemble into a characteristic pinwheel-like organization [24–26], and for maintenance of the epithelial integrity of the ependyma [3, 27, 28]. Given this broad array of functions, abnormalities of the ependymal cells lead to a variety of neurological conditions. For instance, in mammals, ciliary defects of ependymal cells are commonly associated with ventricular defects and hydrocephalus [3, 12, 29–33]. Beside this, motile cilia in the zebrafish have been shown to regulate spine morphogenesis [34] through the formation of a glycoprotein filament in the CSF called Reissner fiber [35, 36]. To further understand how motile cilia instruct brain and spine development across species, it is now crucial to fully resolve the cellular and functional diversity of ciliated cells in the ependyma and the genetic programs driving their differentiation.

In this study, we have determined the ontogeny of ependymal MCCs within the forebrain of the zebrafish. We observed that MCCs emerge at the juvenile stage, co-exist with monociliated cells, and diversify into distinct lineages through development. We dissected the transcriptional regulatory pathways directing the differentiation program of these motile ciliated cells, and found that this involves the complementary and sequential activation of the master regulatory genes of motile ciliogenesis, *foxj1a* and *foxj1b*, and of multiciliation, *gmnc*. Finally, we characterized the role of individual motile ciliated cell lineages in the morphogenesis of the brain and body axis. We show that while critical for proper ventricular development, ciliated cells regulated by *foxj1b* and *gmnc* are largely dispensable for brain and axial morphogenesis. Altogether, our study reveals the diversity of motile ciliated cell-types within the zebrafish forebrain from a molecular, cellular and functional point of view, and uncovers a remarkable degree of similarity as well as differences with ependymal cells of mammals.

## RESULTS

### Glutamylated tubulin in ciliary axonemes is a reliable marker for *foxj1*-expressing cells

Previous work had suggested that glutamylated tubulin staining could be a specific marker of motile cilia in zebrafish [37, 38]. Indeed, we had observed that ciliary tubulin glutamylation was increased in a subset of cells in the nervous system, which expressed the zebrafish *Foxj1* orthologs*, foxj1a* or *foxj1b* [14, 23]. To further scrutinize the causality between *foxj1* expression and axonemal tubulin glutamylation, and hence its validity as motile cilia marker, we overexpressed Foxj1a in zebrafish embryos using a heat-shock inducible *foxj1a* transgene [39, 40], and monitored the levels and distribution of glutamylated tubulin-positive cilia. We have previously shown that overexpression of Foxj1a using this strategy is sufficient to induce ectopic motile cilia formation in tissues that normally differentiate immotile primary cilia, such as the trunk musculature and eyes [39, 40]. We observed that the short primary cilia formed in these tissues under control conditions have a very low to undetectable levels of ciliary glutamylated tubulin. Strikingly, staining of heat-shocked versus control animals revealed that transgenic overexpression of *foxj1a* was sufficient to induce circa 1000-fold higher glutamylated tubulin levels in cilia in the trunk and eye regions (Supplemental Figure S1A,B,F), in addition to significantly increasing ciliary length (Figure S1A,C,G). We also observed that, in comparison to acetylated tubulin, glutamylated tubulin is not uniformly distributed across the cilium but is enriched at one end of the axoneme (Figure S1D,E,H,I,J), corresponding to the ciliary base marked by gamma-tubulin (Figure S1K) [38, 41, 42]. Taken together, these results revealed that the presence of glutamylated tubulin is a reliable marker for cilia of *foxj1*-expressing cells in the zebrafish.

### Motile ciliated cell abundance and onset of multiciliation correlate with expansion of brain ventricles and parenchyma during development

Having identified a reliable marker for motile cilia, we next tracked the appearance of such cilia and the transition from monociliated to MCCs over the course of forebrain development. We focused on larval (4, 14 days post fertilization (dpf)), juvenile (28-32 dpf) and adult stages, and visualized the presence of cilia by immunostaining and ventricular size upon microinjection of a fluorescent dye.

We observed that concomitant with brain volume expansion from 4 to 14 dpf (Figure 1A1,A2), more cilia appeared on the dorsal telencephalon anterior to the forebrain choroid plexus (ChP) (Figure 1B,C). At 28-32dpf, when major cognitive, learning, and social skills are acquired [43–49], we found that ciliated cells on the dorsal telencephalon increased not only in number, but also started to harbor brushes of cilia reminiscent of MCCs (Figure 1A3,D1,D2). At this stage, MCCs were present on the dorsal telencephalon (Figure 1D2) and in the forebrain ChP (Figure 1D4), and co-existed with monociliated cells (arrows versus arrowheads). Interestingly, the MCCs were not randomly distributed, but were enriched at the midline of the dorsal telencephalon (Figure 1D2, quantified in Figure 1D3) and anterior part of the forebrain ChP (Figure 1D4). At later developmental stages (Figure 1A4,E1), MCCs covered large parts of the dorsal telencephalon, yet, remained enriched around the midline. Staining with a membrane marker (beta-catenin) revealed that cilia do not populate the entire apical surface of these cells uniformly, but are translationally polarized (Figure 1E2), resembling mammalian ependymal MCCs [50, 51]. The adult forebrain ChP, which consists of multiple connected cavities (Figure 1E3), was also composed of large numbers of MCCs, closely apposed to monociliated cells (Figure 1E4), with both cell-types directing their cilia towards the lumen of the cavities.

**Figure 1:**
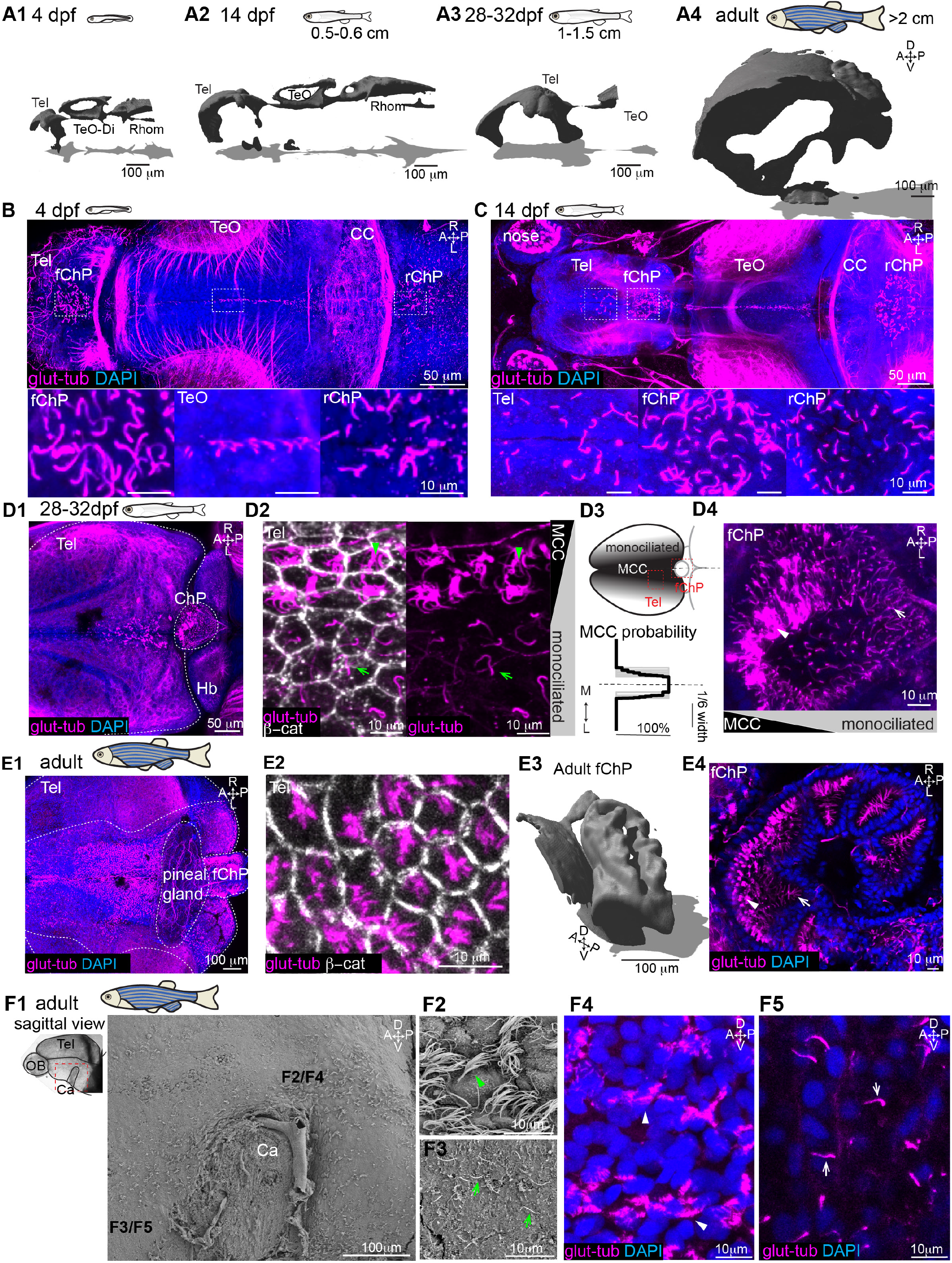
Multiciliation and ventricular/parenchymal expansion correlate during development. **(A1-A4)** Brain ventricles expand from early larval stage (4 dpf, **A1**, n=4), to late larval (14 dpf, **A2**, n=4, 0.5-0.6 cm), juvenile (28-32 dpf, 1-1.5cm long, **A3**, n=3) and adult stages (older than 2 month, larger than 2 cm, **A4**, n=3) as shown upon 3D reconstruction of brain ventricles injected with RITC-dextran and imaged by confocal microscopy. **(B)** At 4 dpf, single glutamylated tubulin-positive cilia are located on the larval forebrain ChP (fChP), on the dorsal roof and ventral part of the tectal/diencephalic ventricle and in the rhombencephalic ChP (rChP). (n=5). **(C)** At 14dpf, cilia number increases along the dorsal telencephalon, rostral to the fChP and in the rChP. Cells are monociliated throughout the brain (n=3). **(D1)** At 28-32 dpf, brushes of glutamylated tubulin-positive cilia appear on the dorsal telencephalon (n=3). **(D2)** Cilia belong both to monociliated cells (green arrow) and MCCs (green arrowhead) as shown upon co-staining with the membrane marker beta-catenin. **(D3)** MCCs are located medially as compared to monociliated cells (quantified in bottom panel, n=9). **(D4)** The fChP at 28-32dpf is composed of mono- and MCCs, which are arranged in an anterior-posterior manner (n=3).**(E1)** In the adult brain, MCCs are enriched in the medial part of the dorsal telencephalon. **(E2)** Cilia do not cover the entire apical surface of MCCs as shown upon co-staining with beta-catenin (n=4). **(E3)** Adult ChP folds into multiple interconnected cavities, as shown upon injection of RITC dextran and 3D reconstruction of confocal images (n=4), and remain composed of mono- and MCCs **(E4)** (n=3). **(F1)** SEM analysis of the midline of an adult telencephalon (n=3) in the region surrounding the anterior commissure (Ca) highlighted in red, reveals the presence of MCCs (**F2)** and monociliated cells (**F3**). Location of F2-F5 is indicated in F1. **(F4-F5)** Confocal image of the midline of the telencephalic hemisphere immunostaining with glutamylated tubulin shows MCCs **(F4)** and monociliated cells **(F5)**. A: anterior, P: posterior, D: dorsal, V: ventral, M: medial, L: lateral, Tel: telencephalon, TeO: optic tectum, Rhomb: rhombencephalon, CC: cerebellum, Hb: habenula, MCC: multiciliated cells.

Next, using scanning electron microscopy (SEM) and glutamylated tubulin staining we observed that ciliated cells were also present along the ventral part of the telencephalon, on the midline surrounding the anterior commissure (Figure 1F1-F2). Ciliated cells carried either a ciliary brush (Figure 1F4) or a single cilium (Figure 1F5), similar to the dorsal telencephalon.

Altogether, our observations show that as the brain parenchyma and ventricular cavities expanded in size, the numbers of motile ciliated cells increased and the multiciliation program was initiated.

### All ciliated cells on the dorsal telencephalon and ChP express *foxj1b*, while a subset on the ventral telencephalon express *foxj1b*

To elucidate the genetic identity of the motile ciliated cells, we analyzed the expression of the master regulator of motile ciliogenesis, *Foxj1*. The two zebrafish *Foxj1* orthologs, *foxj1a* and *foxj1b*, are differentially expressed in the larval brain [14]. Since we previously identified that ciliated cells in the telencephalon specifically express foxj1b, we monitored the expression of *foxj1b* through development. To this end, we employed the GFP-reporter line *Gt(Foxj1b:GFP)^tsu10Gt^*, which contains GFP inserted within the coding sequence of the *foxj1b* gene and faithfully labels motile ciliated cells in the larval brain [14]. From 4 dpf to adult stage, we detected a gradual expansion of *foxj1b*-expressing cells along the anterior and lateral part of the dorso-medial telencephalon and in the ChP (Figure 2A, 2B, 2C1, 2D1). All *foxj1b:GFP*-expressing cells in the telencephalon and ChP harbored glutamylated tubulin-positive cilia and *vice versa* (Figure 2A,B,C1,D2,D3), implying that *foxj1b* plays a major role in these cells. Interestingly, at 1 month of age, we noted that cells located at the midline of the telencephalon not only harbored more cilia per cell, but also expressed *foxj1b:GFP* more intensely (Figure 2C2-C4), suggesting that *foxj1b* levels and activity of the multiciliation program might be correlated.

**Figure 2:**
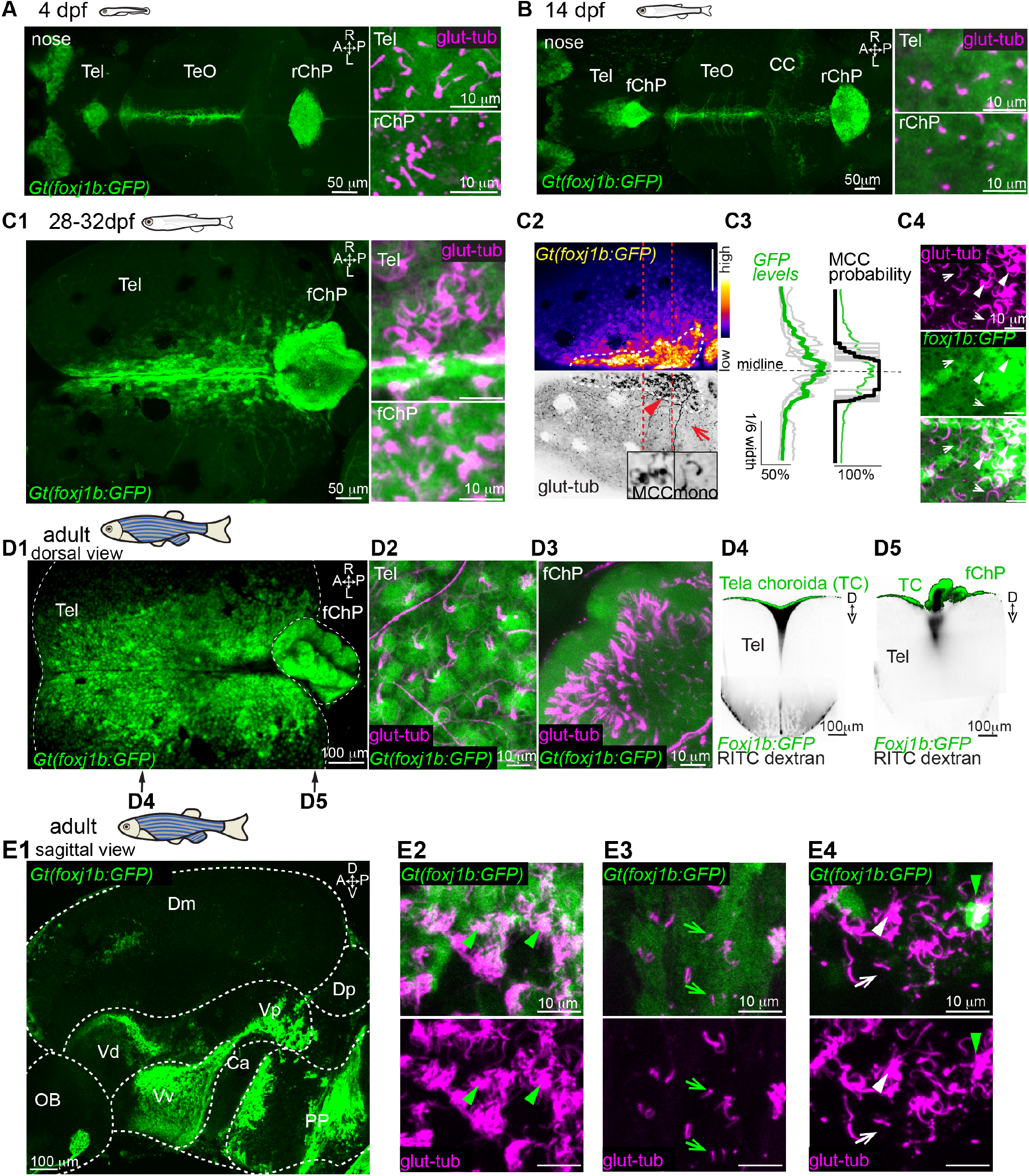
Ciliated cells located on the tela choroida (TC), telencephalon and ChP express foxj1b to a different extent. **(A-D)** Confocal images of Gt(foxj1b:GFP)^tsu10Gt^ transgenic line immunostained for glutamylated tubulin (magenta) at 4 dpf (n=4) **(A)**, 14 dpf (n=3) **(B)**, 28-32 dpf (n=4) **(C)** or adult stages (n=3) **(D)**. **(A)** At 4 dpf and **(B)** 14dpf, foxj1b:GFP expressing cells are located in the dorsal telencephalon, dorsal roof of the optic tectum and rChP and are ciliated. **(C1-C4)** At 28-32 dpf, foxj1b:GFP expressing cells are located on the medial part of the telencephalon and on the forebrain ChP (fChP) **(C1)**. **(C2)** GFP levels are higher in cells located in the medial part of the dorsal telencephalon (indicated with a dashed white line), which is enriched in MCCs. The arrowhead shows a MCC and the arrow shows a monociliated cell, also magnified in the inset. **(C3)** Quantification showing higher GFP levels (relative GFP levels, n=6) and multiciliation (probability of n=9, as shown in Figure 1D3) in the medial telencephalon (region indicated with a red line in panel C2). **(C4)** MCCs (arrowhead) express higher foxj1b than neighboring monociliated cells (arrow). **(D1-D5)** At adult stages, **(D1)** foxj1b:GFP expressing cells are located primarily in the medial part of the dorsal telencephalon **(D2)** and choroid plexus **(D3)** and are ciliated. **(D4-D5)** foxj1b:GFP expressing cells are in direct contact with CSF and are present **(D4)** on the TC, which is the epithelial layer located above the CSF-filled cavity, and **(D5)** on cells in the adult forebrain ChP surrounding CSF filled cavities, as shown upon injection of RITC-dextran (black) in a Gt(foxj1b:GFP)^tsu10Gt^ brain explant. Location of the optical section of D4 and D5 are indicated in D1 (n=3). **(E1)** Confocal image of the midline of the telencephalic hemisphere of a Gt(foxj1b:GFP)^tsu10Gt^ transgenic animal revealed that foxj1b:GFP expressing cells are located mostly in the ventral telencephalon and diencephalon (OB: olfactory bulb, Dm: dorso-medial, Dp: dorso-posterior, Vd: ventro-dorsal, Vp: ventro posterior, Vv: ventro-ventral, PP: Preoptic nucleus, Ca: Anterior commissure). (n=3). **(E2-E4)** Immunostaining with glutamylated tubulin shows the foxj1b:GFP expressing cells harbor multiple cilia (green arrowhead in **E2** and **E4**) or only one cilium (green arrow in **E3**). **(E4)** Some ciliated cells with multiple cilia (white arrowhead) or a single cilium (white arrow) are GFP negative. A: anterior, P: posterior, D: dorsal, V: ventral, M: medial, L: lateral, Tel: telencephalon, TeO: optic tectum, Rhomb: rhombencephalon, CC: cerebellum, Hb: habenula, MCC: multiciliated cells.

To ascertain where the ciliated cells are in relation to the brain parenchyma, we injected a fluorescent dye within the ventricles of brain explants from *Gt(Foxj1b:GFP) ^tsu10Gt^* transgenic animals. We observed that *foxj1b:GFP* expressing cells are not located on the brain parenchyma, but rather in the epithelial layer above the ventricle, known as the tela choroida (TC) [17, 52, 53] (Figure 2D4). We also observed that *foxj1b:GFP*-expressing cells in the ChP surrounded the CSF-filled cavity (Figure 2D5). To discern whether ciliated cells within the telencephalic midline expressed *foxj1b* similar to the dorsal telencephalon, we imaged the midline of the telencephalic parenchyma of Gt(Foxj1b:GFP)*^tsu10Gt^* transgenic animals. We observed that *foxj1b:GFP*-expressing cells were located mostly on the ventral part of the telencephalon and in the preoptic nucleus of the diencephalon, in the region where we observed ciliated cells by SEM (Figure 1F1). Upon staining with glutamylated tubulin antibody, we identified that *foxj1b:GFP*-positive cells are either non-ciliated, mono- ciliated, or multiciliated (Figure 2E2-E3), and that a proportion of mono- and MCCs were not *fox1b:GFP*-positive (Figure 2E4), suggesting that these cells might express the paralogous gene *foxj1a*.

In sum, our results show that motile ciliated cells express *foxj1b* differentially depending on their location within the telencephalon, thereby inferring that ependymal cells have diverse identities.

### Ciliated cells in the telencephalon and ChP are motile and generate directional CSF flow

To confirm that glutamylated tubulin positive cilia are motile, we performed video microscopy on adult brain explants using transmission light microscopy, followed by a Fourier based analysis [8, 14]. We observed that cilia in the ChP (Figure 3 B1,B2), TC (Figure 3 C1-C2) and ventral telencephalon (Figure 3 D1-D2) beat with frequency ranging from 5 to 35 Hz. Although beat frequencies largely varied, ciliary beating was sufficient to elicit directional fluid flow revealed by particle tracking of injected fluorescent beads (Figure 3E-H). In particular, we observed a clear directionality in the dorsal telencephalic ventricle, with flow from anterior to posterior in the superficial part (Figure 3G1) and posterior to anterior in the deeper region (Figure 3G2). In the ventral telencephalon, we also observed that flow is directionally organized, going rostrally anterior to the commissure and caudally posterior to the commissure (Figure 3H). Thus, ciliated cells identified using glutamylated tubulin staining and *foxj1* expression are motile and generate directional fluid flow.

**Figure 3:**
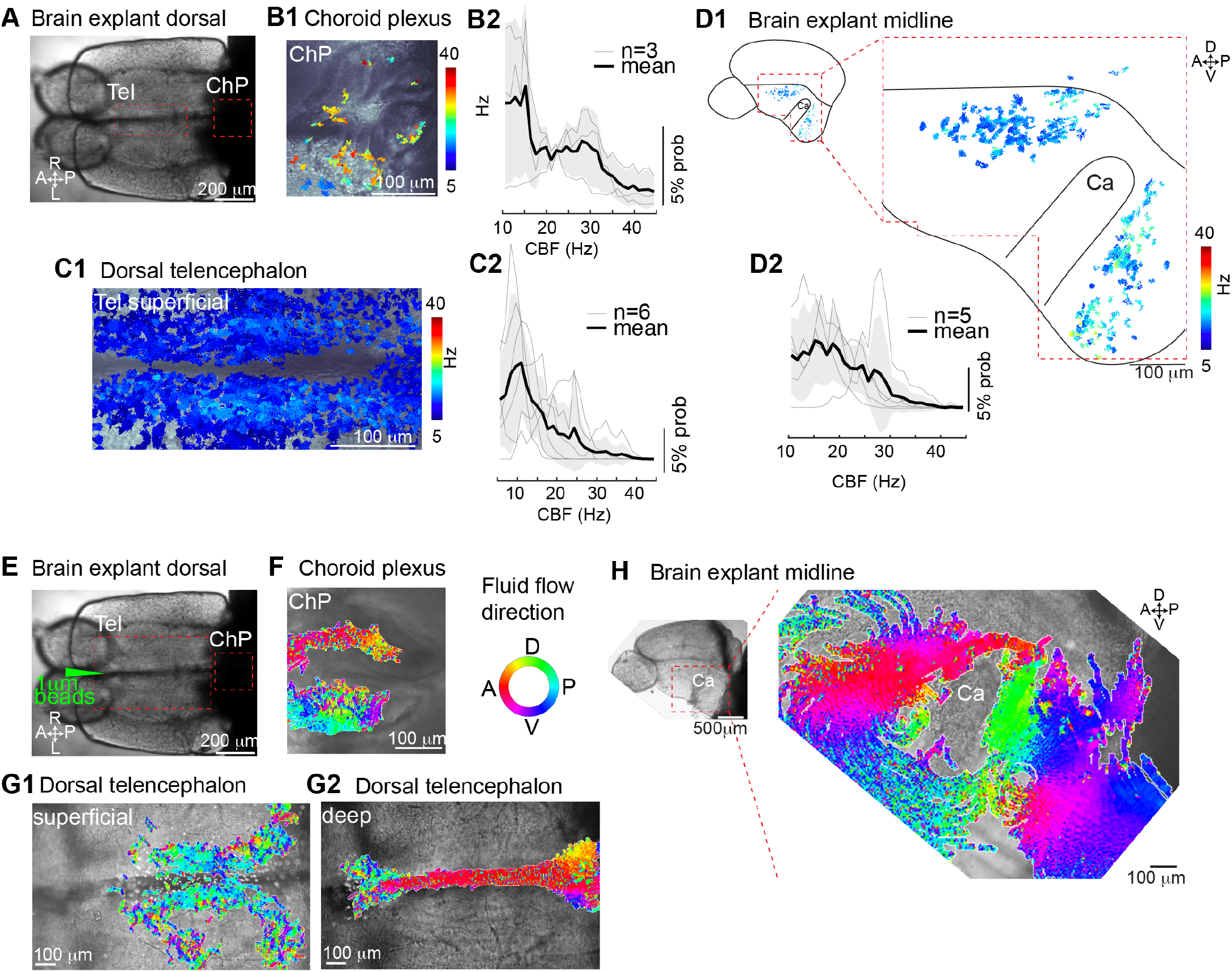
Ciliated cells in the telencephalon and ChP are motile and generate CSF flow. **(A)** Dorsal view of an adult brain explant used for measuring ciliary beating. (**B-C**) Cilia in the forebrain ChP **(B1-B2)** and dorsal telencephalon **(C1-C2)** actively beat as shown with high speed video microscopy followed by a pixel-based Fourier analysis. **(B1)** Map of ciliary beating in the ChP. Ciliary beat frequency (CBF) is color coded. **(B2)** Probability histogram showing the frequency of the pixel-based analysis for n=3 brain explants and the average +/− sem. **(C1)** Map of ciliary beating along the dorsal telencephalon. **(C2)** Probability histogram showing the frequency of the pixel-based analysis for n=6 brain explants and the average +/− sem. **(D1-D2)** Analysis of ciliary beating on a telencephalic hemisphere explant. **(D1)** Ciliary beating is present both anterior and posterior to the anterior commissure (Ca). **(D2)** Probability histogram showing the frequency of the pixel-based analysis for n=5 brain explants and the average +/− sem. **(E)** Dorsal view of an adult brain explant used for measuring fluid flow upon injection of 1 micron fluorescent beads in the anterior part of the dorsal telencephalon. **(F-G)** Directional fluid flow in the choroid plexus **(F)** and dorsal telencephalon **(G1-G2**). Directionality, recovered from single particle tracking, is color-coded. **(F)** In the ChP, fluid flow occurs throughout the cavities (n=4). Note: The directionality of fluid flow is variable and depends on the location within the organ. **(G1-G2)** In the dorsal telencephalon, fluid flow is directed **(G1)** caudally in the superficial part (n=5) and **(G2)** rostrally in the deeper part (n=5). **(H)** In the telencephalic midline, fluid flow is directed rostrally anterior to the anterior commissure and caudally posterior to the anterior commissure (representative example of n=5, **E2**).Tel: telencephalon, ChP: choroid plexus, A: anterior, P: posterior, D: dorsal, V: ventral, R: right, L: left,

### Multiciliation is driven by a genetic program involving *gmnc*

We next sought to identify genes responsible for the transition from mono- to multiciliation. Two genes have been previously reported to transcriptionally regulate multiciliation -*Mcidas* [54, 55] and *Gemc1/Gmnc* [56–60]. Since *gmnc* plays a more dominant role than *mcidas* in zebrafish [59, 61, 62], we chose to focus on *gmnc* mutants. We analyzed *gmnc* mutants at 1 month (Figure 4A), when multiciliation starts to occur, as well as in adult stages (Figure 4B). We observed that ciliated cells in all parts of the telencephalon and ChP of *gmnc* mutants did not harbor ciliary brushes but instead single cilia. Similar results were obtained by SEM analysis of the rhombencephalic ventricle (Figure 4C). Since ciliated cells in *gmnc* mutants retained a glutamylated tubulin-positive solitary cilium, we examined whether they maintained the expression of *foxj1b*. We imaged the brain of 1 month old *gmnc* mutants that also carried the *Gt(Foxj1b:GFP)^tsu10Gt^* transgene. We observed that *foxj1b* remained expressed in the ciliated cells despite the absence of *gmnc* activity (Figure 4D). These findings establish that *gmnc* is essential for inducing multiciliation in motile ciliated cells of the telencephalon and ChP, but is not required for motile ciliogenesis *per se*.

**Figure 4:**
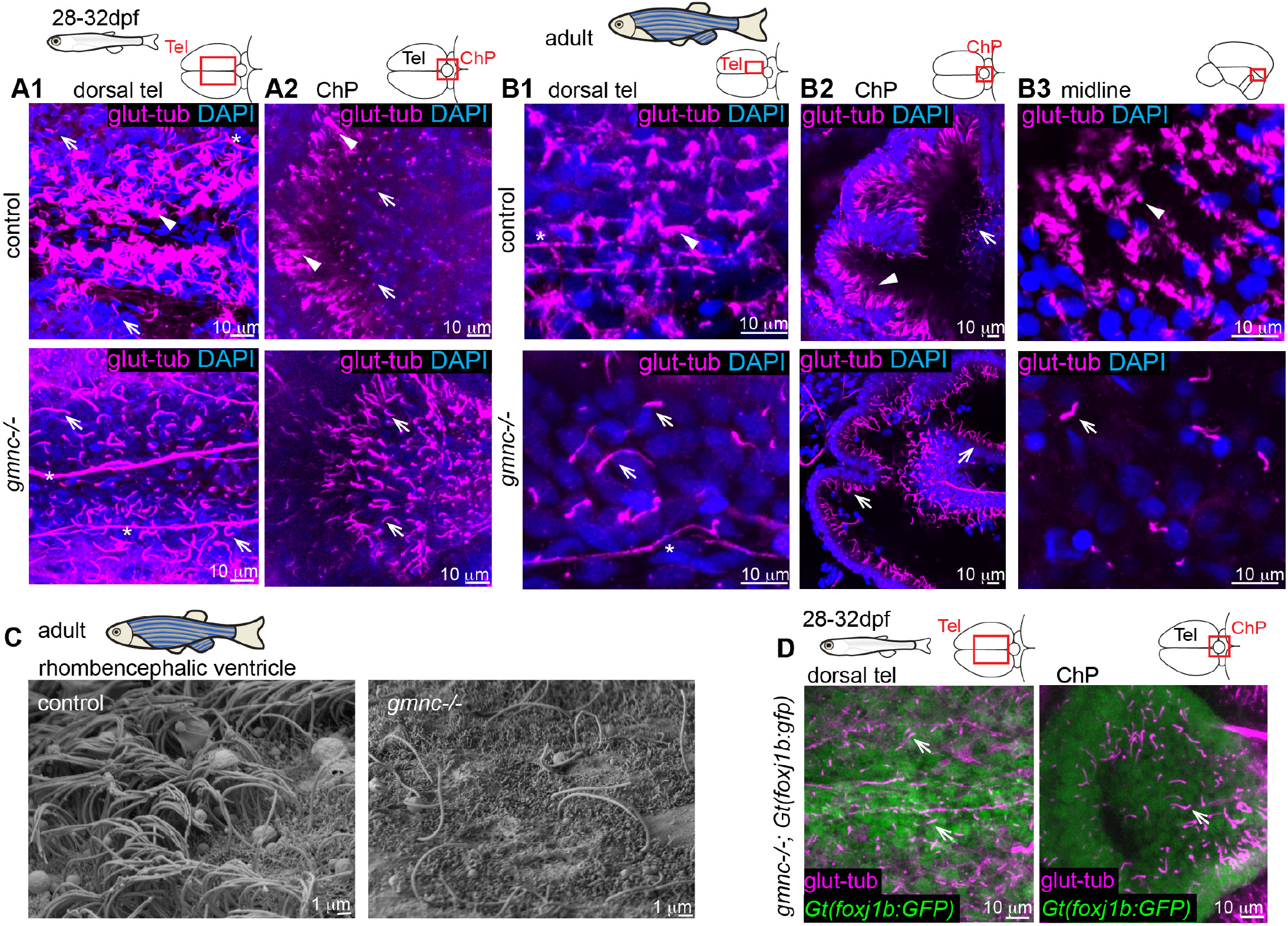
gmnc is required for multiciliation in ependymal cells. **(A-B)** Confocal images of 1 month old **(A1-A2**, n=3 ctrl 3 mutant) and adult brains **(B1-B3**, n=3 ctrl 3 mutant) immunostained for glutamylated tubulin (magenta) and DAPI (blue) revealed the absence of MCCs (arrowhead) in gmnc mutant. All ciliated cells in gmnc mutant are monociliated (arrow). Images were collected as indicated by the red box on the scheme at the top right. * indicate axons. **(A1)** 1 month old dorsal telencephalon (Tel). **(A2)** 1 month old ChP (ChP). **(B1)** Adult dorsal telencephalon. **(B2)** Anterior portion of the adult ChP. **(B3)** Adult midline above the anterior commissure (n=3). **(C)** Scanning electron microscopy images of the rhombencephalic ventricle (n=3). **(D)** Confocal images of a gmnc^−/−^;Gt(foxj1b:GFP)^tsu10Gt^ animal reveal that foxj1b:GFP remained expressed in monociliated cells (arrow) in the dorsal telencephalon (left) and ChP (right) (n=4).

### *foxj1b* induces monociliation in the telencephalon and ChP

Since *foxj1b* is expressed in a large number of ciliated cells in the telencephalon and ChP, we sought to identify the function of *foxj1b* in these cells. For this, we generated a mutant allele, *foxj1b^sq5719^*, lacking most of the coding sequence (see Supplemental Figure S2 and Materials and Methods). Homozygous mutants are viable, and do not exhibit obvious morphological abnormalities other than otolith defects in the embryonic inner ear, consistent with our earlier work using morpholinos [63]. We then investigated the presence of ciliated cells in the brain of these mutants from larval to adult stages. At 4 dpf, we observed that *foxj1b* mutants lacked cilia in the dorsal telencephalon, but not in the cells surrounding the optic tectum (Figure 5A). To identify whether these remaining ciliated cells were dependent on the expression of the paralogue *foxj1a*, we generated *foxj1a/foxj1b* double mutant larvae using our previously established *foxj1a* mutant strain [14]. We observed a complete loss of cilia from the brains of double *foxj1a/b* mutant larvae (Figure 5A bottom panel), confirming that formation of glutamylated tubulin-positive cilia is instructed by different combinations of *foxj1* genes. We next analyzed the brains of 2 week old *foxj1b* mutants (Supplemental Figure S3A), prior to multiciliation, and observed that all cilia in the dorsal telencephalon were absent, as seen in 4 dpf larvae. Surprisingly, at one month of age, *foxj1b* mutants harbored similar numbers of MCCs (arrowhead) as control animals, both in the dorsal telencephalon and ChP (Figure 5 B1-B2). At this stage, we observed that monociliated cells (arrow) remained absent in the lateral part of the dorsal telencephalon (inset in Figure 5B1) and in the posterior part of the forebrain ChP (Figure 5B2, indicated by #). Further analyses of the adult brain by confocal (Figure 5C) and SEM analysis (Supplemental Figure S3B) revealed that MCCs were not particularly affected in *foxj1b* mutants. However, monociliated cells, in particular those in the ChP, were absent even at adult stages (Figure 5C2). Altogether, these results indicate that *foxj1b* is required for the formation of monociliated cells in the dorsal telencephalon and ChP, but is not essential for MCC differentiation.

**Figure 5:**
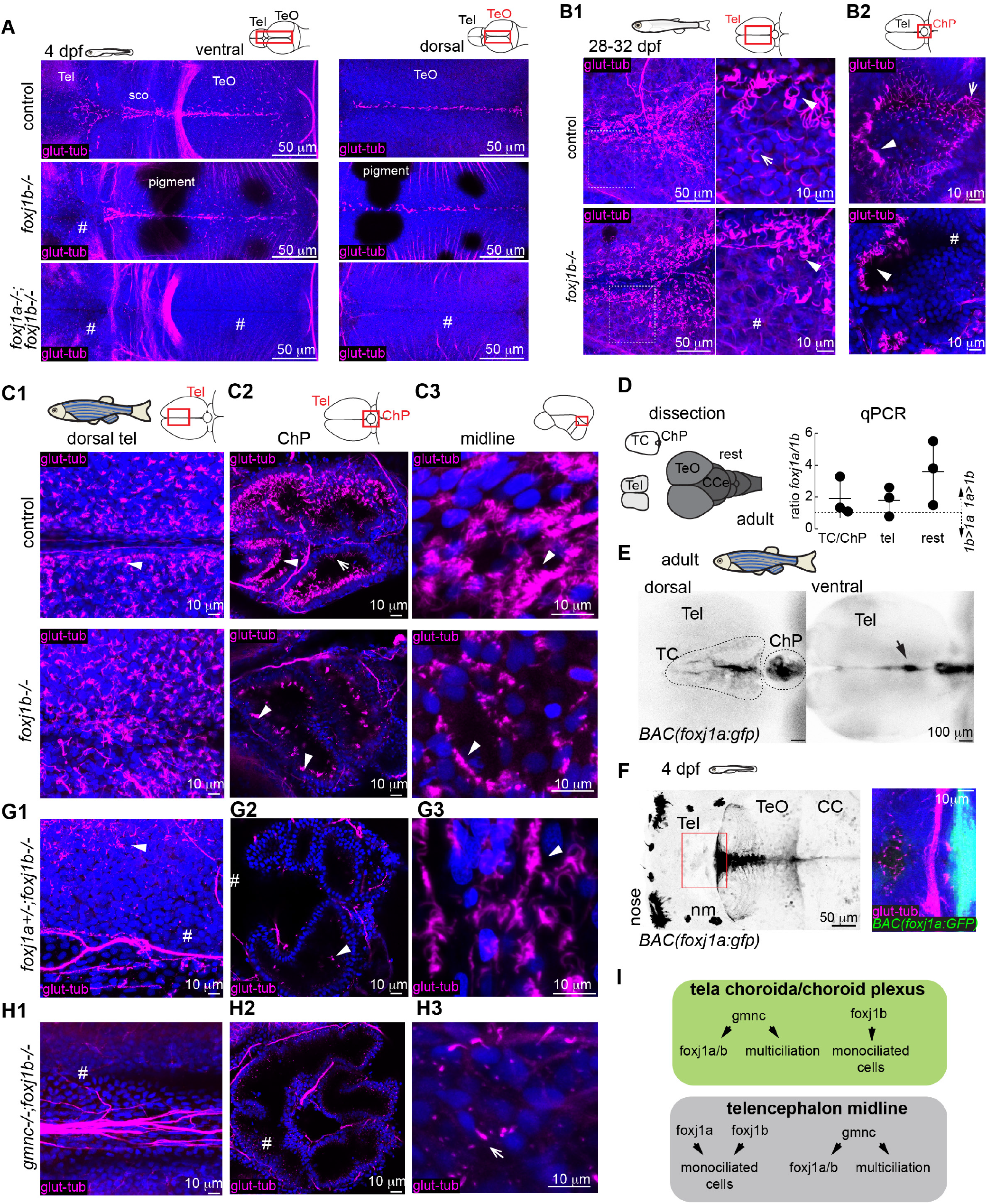
foxj1a and foxj1b play diverse roles in the formation of motile ciliated cells. **(A)** Confocal images of 4 dpf larvae immunostained for glutamylated tubulin (magenta) and DAPI (blue) showing lack of cilia in the dorsal telencephalon in foxj1b mutant (middle, n=9) and throughout the larval ventricular system in double foxj1a/b mutant (n=6). Cilia loss is indicated by #. Maximum projection at two different depths are shown. **(B1-B2)** Confocal images at 1 month of age shows that MCCs (arrowhead) are not affected in the foxj1b mutant on the dorsal telencephalon **(B1**, n=6) and ChP **(B2**, n=6). Cilia remain absent in the monociliated cells (arrow) located laterally to the MCCs as shown in the magnified inset. Cilia loss is indicated by #. **(C1-C3)** Confocal images show the MCCs are not affected in the adult dorsal telencephalon **(C1**, n=3) and remain present in the choroid plexus **(C2**, n=4). **(C3)** Ciliated cells remain in the midline of the telencephalon/diencephalon of foxj1b mutants (n=3). **(D)** RT-qPCR analysis on dissected TC/ChP, whole telencephalon and the rest of the brain shows that foxj1a is expressed in the adult TC/ChP (n=3). **(E-F)** foxj1a is expressed in the dorsal telencephalon, ChP and telencephalic midline of the adult brain **(E)** but not in the larval 4dpf brain **(F)** as shown using a BAC transgenic line Tg(foxj1a:gfp). **(G1-G3)** Confocal staining of an adult foxj1a+/−;foxj1b−/− double mutant immunostained for glutamylated tubulin (magenta) and DAPI (blue) shows a partial loss of MCCs (indicated by #) in the TC (**G1)** and a major loss of cilia in the ChP (**G2**). (**G3**) MCCs (indicated by arrowhead) remained in the telencephalic midline of foxj1a+/−;foxj1b−/− (n=5). **(H1-H3)** Confocal staining of an adult gmnc;foxj1b double mutant immunostained for glutamylated tubulin (magenta) and DAPI (blue) show a complete loss of cilia in the TC and ChP (**G1** and **G2**). **(G3)** Single cilia remained in the telencephalic midline of double mutants (indicated by an arrow) (n=4). (**I)** Diverse genetic programs instruct the formation of ciliated cells in the TC/ChP and in the telencephalon midline. nm: neuromast, CCe: cerebellum, TeO: optic tectum, sco: sub-commissural organ.

### *foxj1a* and *foxj1b* play overlapping roles in MCC formation

Since *foxj1b* mutation leads to ciliary defects in mono- but not MCCs, we hypothesized that *foxj1a* expression might be induced upon multiciliation, and thus, be sufficient for MCC formation. To test this, we first examined whether *foxj1a* is expressed in the TC and ChP. We dissected the TC/ChP from the whole telencephalon and the remainder of the brain, and performed qRT-PCR (Figure 5D). Using this approach, we confirmed that *foxj1a* is expressed in the adult TC/ChP. Our results also indicated that *foxj1a* expression is slightly higher than *foxj1b* throughout the adult brain (Figure 5D). Therefore, *foxj1a* expression might be sufficient to promote the formation of MCCs in *foxj1b* mutants. To identify the location of *foxj1a*-expressing cells on the TC, we generated a novel bacterial artificial chromosome (BAC)-based reporter transgenic strain. Analysis of this strain confirmed the expression of *foxj1a* in the TC and ChP in 2 month-old adults, in addition to the telencephalic midline (Figure 5E). As we did not observe *foxj1a* expression in the larval ChP (Figure 5F), our results suggest that *foxj1a* is expressed when ciliated cells acquire multiciliation. To test the impact of *foxj1a* in MCC ciliation, we generated *foxj1a+/−;foxj1b−/−* mutants. In contrast to *foxj1a−/−*, which are embryonic lethal due to severe body curvature [14, 34] we were able to identify *foxj1a+/−;foxj1b−/−* viable adults. In the brain of these mutants, we observed reduced numbers of cilia in the ChP compared to wild-type control or *foxj1b*−/− (Figure 5G2). In the TC, we also observed reduced numbers of MCCs, but with a variable degree of penetrance (Figure 5G1 and S4). However, large numbers of MCCs remained in the telencephalic/diencephalic midline (Figure 5G3). These results highlight that *foxj1a* and *foxj1b* play important and overlapping roles for MCC ciliation in the telencephalon, and that one copy of *foxj1a* gene is sufficient for MCC differentiation in the TC and midline.

To test whether *foxj1a* activity depends on the expression of *gmnc* as previously shown [54, 56–59, 61], we generated *gmnc/foxj1b* double mutants. In these animals, which are viable, we observed cilia loss on the TC and ChP (Figure 5H1-H2). By contrast, solitary cilia persisted in the telencephalic midline (Figure 5H3), most likely associated with the cells that exclusively express *foxj1a*. In sum, these results highlight that the diversity of motile ciliated cells within the zebrafish brain ventricles expands during development, and depends on the sequential or parallel activation of *foxj1* and *gmnc*-dependent transcriptional programs (Figure 5I).

### *foxj1a* and *foxj1b* are expressed in two ependymal cell lineages and in different subsets of neuronal progenitors

To further unravel the genetic diversity of ependymal lineages, we performed single cell transcriptomic analysis of an adult telencephalon, using a protocol as described previously [64]. We labelled cells located in close proximity to the ventricle by injecting a fluorescent cell tracer in 1-year old animals [65]. We then FACS sorted telencephalic cells, performed single-cell sequencing and recovered 3158 single cells following filtering (see Methods for details and Figure S5A). After multiple steps of clustering, we identified various cell-types including neurons, immune cells, oligodendrocytes, quiescent progenitors and more actively proliferating neuroblasts (Figure 6A1 and S5). To identify potential multiciliated ependymal clusters, we searched for cells expressing *foxj1a* and *foxj1b (*Figure 6A2), *gmnc* (Figure 6A3), and previously reported ependymal markers from mice [66] and zebrafish [64] including *enkur* ([67] Figure 6A4) and *mia* (Figure 6A5). Based on these criteria, we identified two potential clusters of ependymal-like cells: the PC cluster 14 and the *dkk* cluster (Figure 6A,B). Interestingly, one of these ependymal clusters, the PC cluster 14, is highly similar to progenitor-like cells, but not the *dkk* cluster. We also identified that only the *dkk* cluster expressed the ependymal cell marker *mia* (Figure 6A5, B5). Strikingly, we observed that *foxj1a* and *foxj1b* expression was not limited to these two ependymal cells clusters, but the genes were broadly expressed in progenitor-like cells (Figure 6A2,B2). Most of these cells expressed either *foxj1a* or *foxj1b*, with circa 15% expressing both *foxj1a* and *foxj1b* (Figure 6C). *gmnc* was also expressed in both ependymal-like clusters (Figure 6A3,B3). Since *gmnc* is expressed transiently when the cells initiate multiciliation [56, 58, 59], our results suggest that these two populations of ependymal cells are most probably generated independently, and do not represent cells at different developmental stages. Moreover, we observed that all *gmnc*-expressing cells were either *foxj1a* and/or *foxj1b* positive (Figure 6D), confirming previous observation that *gmnc* induces *foxj1* expression [54, 56–59, 61]. Next, we parsed the differentially regulated genes between these two ependymal-like clusters (Supplemental Figure S5D), and identified a total of 257 genes. Among these, radial glia/astroglia markers (*her4.1,her4.2, gfap* [68]) were enriched in the PC cluster 14, while other genes, including those encoding secretory proteins present in CSF (*rbp4* [69]), marker of ChP (*igfbp2a* [70]), or mammalian ependymal cells (*mia* [66]) were enriched in the *dkk* ependymal cluster (Figure 6A5).

**Figure 6.**
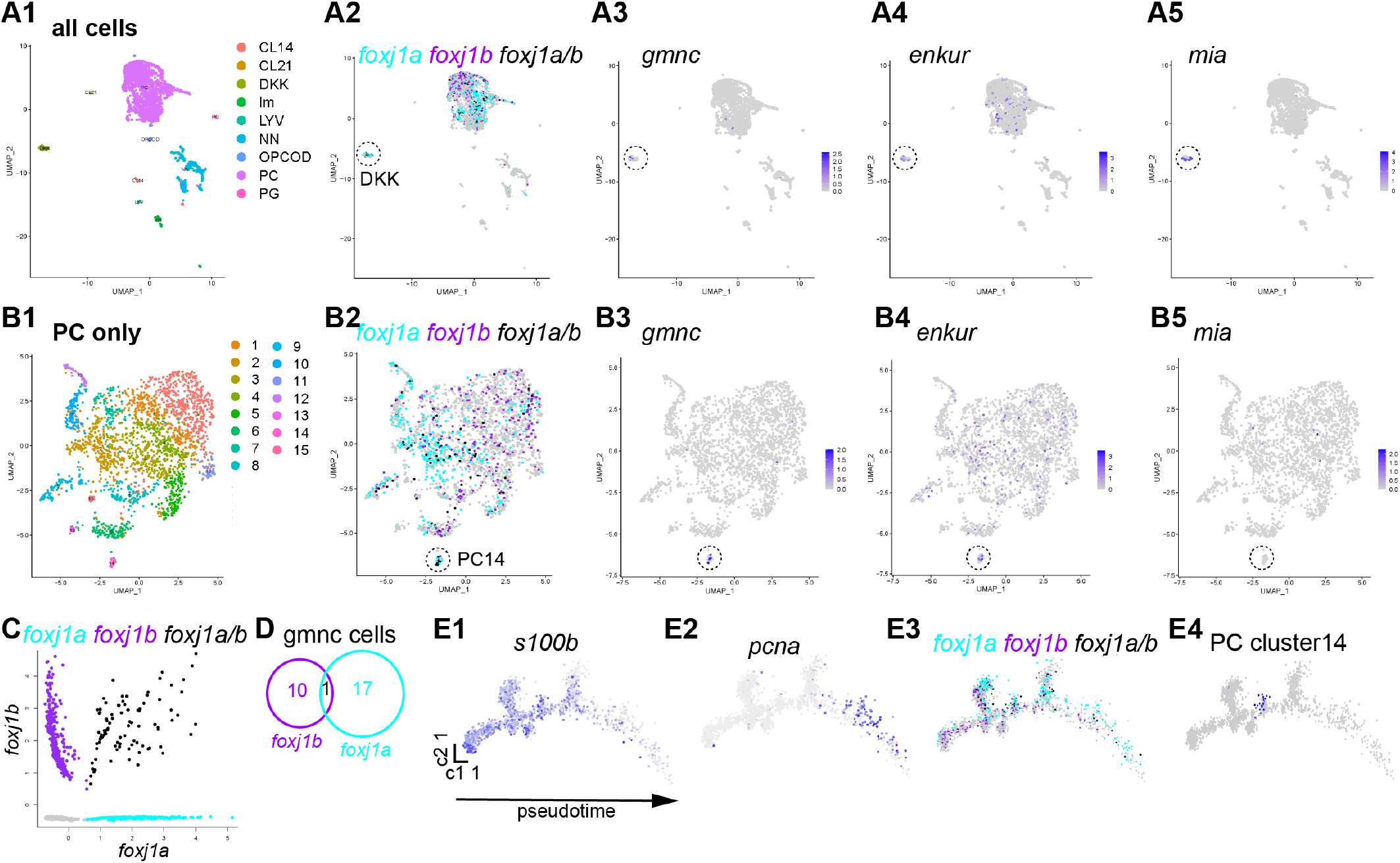
Diversity of motile ciliated cells in the adult telencephalon revealed by single cell RNA sequencing analysis. **(A-B)** Unbiased clustering analysis of all cells **(A1)** or all progenitor-like cells (PC, **B1)** revealed the presence of a large number of cell clusters, including two potential ependymal clusters. Ependymal clusters DKK **(A1-A5)** and PC14 **(B1-B4)** were identified based on the expression of the motile ciliogenic genes foxj1a/b (foxj1a only cells in cyan, foxj1b only cells in magenta, foxj1a/b-co-expressing cells in black **A2**, **B2**) and gmnc (**A3**, **B3**), and ependymal markers enkur (**A4**, **B4**) and mia (**A5**, **B5**). **(C)** Scatter plot representing the expression levels of foxj1a and foxj1b in all cells. (**D)** All cells with gmnc expression (total of 27) also expressed foxj1a and/or foxj1b. **(E)** Pseudotime analysis of the PC cells plotted using Monocle algorithm showed a progression from quiescent (s100b, **E1**) to a proliferative stage (pcna, **E2**). **(E3)** foxj1b is expressed more in quiescent progenitors while foxj1a is expressed more in proliferative progenitors. **(E4)** PC14 ependymal cells (indicated in blue) branch out from quiescent progenitors.

We then sought to unravel the identity of “non-ependymal” *foxj1*-expressing progenitor-like cells. We plotted all progenitor cells on a cell-trajectory using Monocle (Figure 6E1-E4) [71]. Using this approach, we were able to monitor the progression of progenitor cells from a quiescent (shown by expression of e.g. *s100b*, Figure 6E1) to a more proliferative neuroblast state (shown by expression of e.g. *pcna*, Figure 6E2). We observed that *foxj1b* is expressed preferentially in quiescent progenitors, while *foxj1a* is expressed at higher levels in proliferative neuroblasts (Figure 6E3). This analysis also revealed that the “progenitor-like” PC cluster 14 ependymal cell cluster branches out from radial glia-like quiescent progenitors (Figure 6E4), in agreement with similar deductions with the mouse brain [4, 6, 7]. Altogether, our single cell RNA sequencing data have delineated that there are at least two different ependymal cell clusters with different origins, and that *foxj1a* and *foxj1b* are expressed not only in ependymal cells but also in neuronal progenitors.

### Loss of *foxj1b* and *gmnc* does not impact body axis and brain morphogenesis, but affects the size of brain ventricles

Previous work using zebrafish revealed that motile cilia defects induce severe scoliosis of the spine [19] due to aberrations in Reissner fiber formation, impaired CSF flow, catecholamine transport and urotensin II-related peptide gene expression in spinal CSF-contacting neurons [34, 36, 72–74]. To date, it remains unexplored which ciliated cell population contributes to this phenotype. As we did not observe obvious axial malformations among *foxj1b*, *gmnc, foxj1a+/−;foxj1b*−/−, or *foxj1b;gmnc* double mutant adults (Figure 7A), our findings suggest that lack of cilia on *foxj1b*-expressing cells or on MCCs is not the etiological basis of scoliosis.

**Figure 7:**
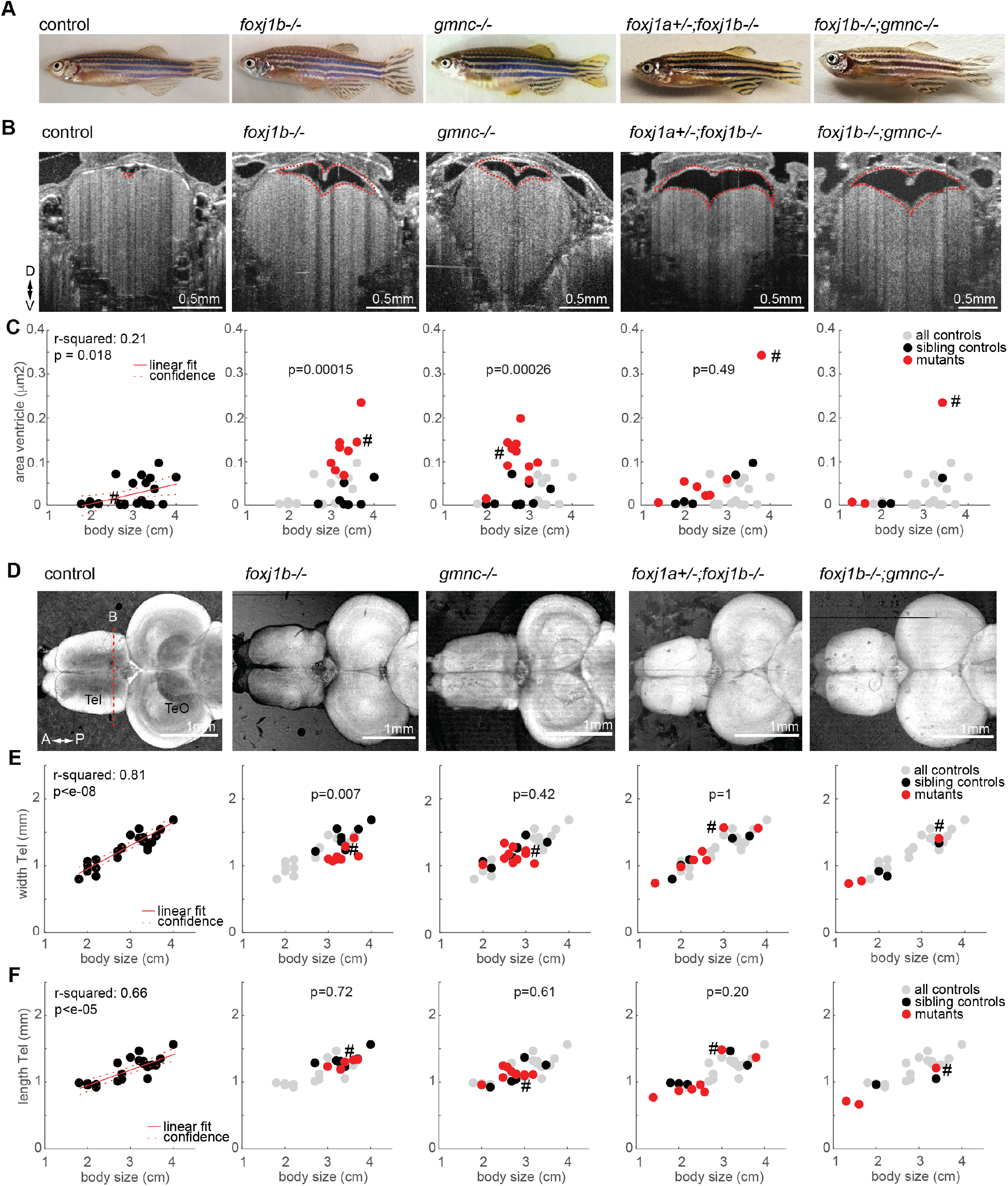
gmnc−/−, foxj1b−/−, foxj1a+/−;foxj1b−/− and gmnc−/−;foxj1b−/− mutants do not display major body and brain malformations, but enlarged telencephalic ventricles. **(A)** Pictures of adult zebrafish showing absence of scoliosis. **(B)** OCT images of anaesthetized adult zebrafish revealed enlarged telencephalic ventricles (highlighted with dashed red line) in animals with motile cilia defects as compared to control. Transverse section taken in the caudal part of the telencephalon as indicated by red dashed line in panel D. **(C)** Quantification of ventricular size in all animals, represented as a function of body length. R-squared indicate the linear relationship between body length and ventricular size as shown in red. Left: all controls pooled, Others: all controls in grey, sibling controls in black, mutants in red. P-value (ranksum between sibling controls indicated in black and red) are indicated in the graph. **(D)** Standard deviation projection of OCT images of adult brain explants showing absence of overt brain malformations in all cilia mutants. Tel: telencephalon, TeO: optic tectum **(E-F)** Quantification of the telencephalic width (E) and length (F), represented as a function of body length. R-squared indicate the linear relationship between body length and brain measurements as shown in red. Left: all controls pooled, Others: all controls in grey, sibling controls in black, mutants in red. Examples shown in B/D are highlighted in the scatter plot in C/E-F with a #. D: dorsal, V: ventral, A: anterior, P: posterior.

Impaired motile cilia are associated with hydrocephalus, both in mice and in human patients [3, 12, 29–33, 75, 76]. To test for such a hydrocephalus phenotype, we used Optical Coherent Tomography (OCT) [77, 78] and visualized the ventricular size in anesthetized adult zebrafish *in vivo*, since dissected brain explants often caused the ventricular system to collapse (Figure S6A). In line with data from mammalian studies, we observed that *foxj1b* and *gmnc* mutants have significantly enlarged ventricles. The trend was similar for *foxj1a+/−;foxj1b−/−* mutants and *foxj1b;gmnc* double mutants, although the results for these genotypes were not statistically significant due to limitations in numbers of animals and age distribution (Figure 7B-C). Finally, to assess the impact of impaired motile cilia on overall zebrafish brain development, we imaged adult zebrafish brain explants by OCT. Our morphometric analysis showed no overt brain malformations between motile cilia mutants and control animals (Figure 7D-F and S6E).

Thus, the lack of multiciliation or loss of cilia in specific subpopulations of ependymal cells results in ventricular defects, but not in brain malformation as commonly observed in hydrocephalic mice with cilia defects.

## DISCUSSION

We have characterized the processes regulating the formation of motile ciliated cells in the ependyma of the zebrafish brain, and described their role in CSF flow and ventricular development. We showed that as the brain and its ventricles expand, MCCs appear progressively on the TC, ChP and parenchymal surface in the midline. In addition, our work has revealed extensive heterogeneity in the genetics, development, anatomy and functional properties of the motile ciliated cells. Finally, we showed that this heterogenous population of ciliated cells collectively generate a stereotypical pattern of CSF flow, contributing to proper development of the brain ventricles. We relied on various methods to detect the presence of motile cilia, primarily based on the expression of *foxj1* genes, immunostaining with cilia markers and video microscopy. Importantly, our work has established the causality between *foxj1* activity and the glutamylation of axonemal tubulin, detectable using the GT335 anti-glutamylated tubulin antibody. This discovery has important implications as it allows now the detection of potential motile cilia based primarily on an antibody staining, at least in the zebrafish. However, it is to be noted that not all *foxj1* expressing cells have an actively beating cilium. For example, hair cells of the inner ear and lateral line sense organs have non-motile cilia that are also marked by glutamylated tubulin (Figure S7). Thus, glutamylated tubulin positivity cannot be used solely as a marker of cilia motility. High levels of ciliary glutamylated tubulin most likely arises from enzymes glutamylating tubulin, such as *ttll6*, being targets of *foxj1* [40]. Ttll6 was previously shown to be expressed in mouse ependymal cells [37, 41] and to regulate ciliary beating in zebrafish and mice [37, 41]. Together, these data suggest that glutamylation of ependymal cilia is conserved across species and important for ciliary motility.

In the zebrafish, two paralogous *Foxj1* genes exist, namely *foxj1a* and *foxj1b* [39, 79]. Using a combination of mutants and transgenic lines described here and in our previous study [14], we showed that in the larval brain, there are three cell populations that express different combinations of these genes. This diversity further expands as the animals develop. For instance, in the TC, the two *Foxj1* orthologs are expressed at different development stages, with *foxj1b* from early development and *foxj1a* later in MCCs. This raises the question as to why *foxj1a* is needed later in development in this population of cells. It is possible that *foxj1a* is expressed because Gmnc eventually induces the expression of both *Foxj1* orthologs [54, 56–59, 61, 80], without discriminating between the two target genes. Alternatively, MCCs could require both *foxj1* genes to transcribe enough motile cilia-specific gene transcripts for multiciliation. Since *foxj1b* mutants do not show discernible MCC defects in the TC or the midline telencephalon, it is possible that *foxj1a* plays a more important role in these cells. It is presently unclear why the brain requires such a diversity of ciliated cells and whether the two *Foxj1* orthologs are contributing to such diversity. Our previous work revealed that both *foxj1* genes have the same efficacy at generating ectopic motile cilia-like cilia when overexpressed [39]. Likewise, we showed here that both genes are equally important for ciliogenesis in monociliated cells where only one paralog is expressed. Nevertheless, it will be interesting to identify whether the two *Foxj1* orthologs have specific transcriptional programs, in addition to a common one, and if their unique targets could contribute to ciliary diversity.

Our single cell transcriptomics analysis revealed that, beyond MCCs, neuronal progenitors also express *foxj1*. We found that *foxj1b* is expressed more in quiescent stem cells, *foxj1a* more in proliferative neuroblasts, and only a small percentage of cells express both genes. This observation correlates with the description that radial glial cells harbor cilia with motile cilia-like features in the adult zebrafish telencephalon [16]. *Foxj1* was also shown to be expressed in neuronal progenitors in the embryonic and adult neurogenic niche of the mouse [24, 81]. Our single cell data have also revealed that some ependymal MCCs (PC cluster 14) are in many aspects similar to progenitor cells. This relates well with data obtained from the adult mouse lateral ventricle, where ependymal cells and quiescent neural stem cells share a large number of markers [66]. Moreover, our analysis revealed that MCCs from one ependymal cluster (PC cluster 14) are most probably derived from quiescent progenitor cells. This is in many ways similar to the mouse, where radial glia act as progenitors of the ependymal cells during embryonic development [4, 6, 7].

We also showed that motile ciliated cells on the dorsal telencephalon are not located in the brain parenchyma, but instead on the TC, which is the epithelial layer located above the telencephalon [17, 82], and in the ChP. Based on our observations, ciliated cells in the TC and ChP rely on the same transcriptional program, suggesting that these two structures may have more features in common. In fact, the TC is a fold of the pia mater that gives rise to the ChP, prior to the formation of the plexus with the blood vasculature [83, 84]. Therefore, it is not surprising that these two structures could rely on the same ciliogenic transcriptional program. While at the larval stage the TC/ ChP comprise a flat monolayer, its posterior part (located above the habenula [85, 86]) folds into a structure composed of several cavities as development progresses, with cells pointing their cilia toward the cavities. Interestingly, as the ChP folds within multiple cavities, there remains a spatial organization of mono- versus MCCs. The inverted configuration of the zebrafish forebrain ChP is different from that of mammals, where ependymal cells point outwards in the ventricles and bathe in CSF [87]. These different configurations may result from the eversion of the neural tube in the zebrafish, in contrast to evagination of the mammalian brain [47], and the necessity to increase the surface area of the ependymal layer with limited space. Similar to the zebrafish, cells of the mammalian ChP are multiciliated [80, 88, 89] and express Foxj1 [80, 90]. Cilia on the mouse ChP have also been shown to be motile, particularly at the perinatal period [89, 91, 92]. Yet, it is currently believed that these cilia do not contribute significantly to CSF flow [92, 93], but rather could be involved in other cellular processes [89, 91].

Upon loss of the central regulator of multiciliation, *gmnc*, we found that zebrafish ependymal MCCs differentiate a single cilium, implying that Gmnc activity does not impact motile ciliogenesis *per se* but only the process of multiciliation. This contrasts with mouse ependymal cells where loss of *Gmnc/Gemc1* completely abolishes ciliogenesis, and prevents the maturation of ependymal cells [57, 58] and MCCs of the ChP [80]. Thus, ciliary defects in the *gmnc* mutant zebrafish brain resembles more closely the phenotype of *Mcidas* knockout mice, where multiciliated precursor cells are still specified, but do not acquire multiciliation and instead generate a single motile-like cilium [55, 80]. These results are also similar to our earlier report of *gmnc* loss-of-function in the zebrafish embryonic kidneys where MCC precursors differentiate with single motile cilia [59]. We argue that these contrasting effects are due to the way ciliogenic transcription factors are deployed during ciliated cell development in the different organisms. In zebrafish embryonic kidneys and brain, MCC fate is instituted by *gmnc* acting on cells that first express *foxj1a* or *foxj1b*, respectively, and have already differentiated a single motile cilium. Gmnc modifies this monociliated program to drive multiciliogenesis, in part by upregulating the expression of the *foxj1* genes, and on the other hand, by activating genes for multiple basal body production. Consequently, on loss of *gmnc*, these cells by default remain monociliated. By contrast, during MCC formation in the mouse, Gmnc/Gemc1 functions at the top of the hierarchy of ciliary transcription factors: induction of Gmnc/Gemc1 expression in turn activates the expression of *Foxj1* and other regulatory genes involved in ciliation and genes involved in multiple basal body production [57, 58, 60]. Consequently, in this context, loss of Gmnc/Gemc1 completely impairs motile ciliogenesis. All of these data illustrate how combining several motile cilia-related transcription factors, and their differential deployment in one or more cascades, can generate a high diversity of ciliated cell types.

Defects in ependymal cells can have dramatic consequences on brain development. In mice, ependymal dysfunction ultimately results in hydrocephalus with enlarged ventricles and thinning of the brain parenchyma [3, 12, 30, 32], typically leading to the premature death. By contrast, in humans, the prevalence of hydrocephalus upon ciliary dysfunction is rather low [12, 30, 31]. Our *foxj1b* and *gmnc* mutant zebrafish show enlarged ventricles but no severe brain malformation. It remains to be understood whether this difference in hydrocephalus prevalence among species relates to differences in susceptibility to stenosis of the aqueduct, in overall CSF dynamics and/or in additional genetic predisposition. Additionally, defects in motile cilia have been associated with scoliotic malformations of the spine in zebrafish, at larval, juvenile and adult stages [19, 34, 35, 72, 94, 95]. We did not observe scoliosis in *foxj1b*, *gmnc*, as well as *foxj1a+/−;foxj1b−/−* and *foxj1b;gmnc* mutant animals, suggesting that body axis formation is not regulated by the ciliary mechanisms described in this work. Spine morphogenesis may instead rely primarily on *foxj1a*. In support of this view, although we report adult *foxja+/−; foxj1b−/−* animals without spine defects, we have found this genotype among embryos with ventrally curved axes from crosses of *foxj1a+/−;foxj1b+/−* double heterozygous fish (data not shown). This implies that loss of one copy of *foxj1a* in *foxj1b* homozygotes can predispose for axial malformation.

In conclusion, our work has identified the major motilecil iated ependymal cell-types of the zebrafish brain longitudinally through the process of development, and has dissected the genetic mechanisms underlying their differentiation. We believe that these data have laid the foundation for more detailed investigations aimed at unravelling how these different ciliated lineages regulate specific aspects of brain development and physiology as well as their contribution to proper morphogenesis of the body axis.

## ACKNOWLEDGEMENTS

We thank M. Wulliman and A. S. Mathuru for constructive suggestions and critical reading of the manuscript, the Research Support Centre A*STAR Microscopy Platform for SEM analysis, CMCB Flow Cytometry core for flow cytometry, DRESDEN-concept Genome Center in TU Dresden for single-cell sequencing and Thorlabs for loaning us the OCT device (Thorlabs 1300nm SD-OCT system TELESTO) used in this manuscript. We thank S. Eggen, V Nguyen, A. Nygaard and the Trondheim fish facility team for their technical support. This work was supported by a NTNU strategy grant (N.J.Y.), RCN FRIPRO grant 314189 (N.J.Y) and 239973 (E.Y), ERC starting grant 335561 (E.Y.), Helse Midt-Norge Samarbeidsorganet grant (E.Y., N.J.Y), a Boehringer Ingelheim Fonds fellowship (C.R., J.N.H), a postdoctoral fellowship from the National Research Foundation (NRF) of Singapore (Q.T.), a Singapore International Graduate Award (SINGA) (D.R.), funds from the Agency for Science, Technology and Research (A*STAR) of Singapore (S.R.) and German Center for Neurodegenerative Diseases (DZNE) within Helmholtz Association (C.K.). Work in the Yaksi laboratory is supported by NTNU and Kavli Foundation.

## AUTHOR CONTRIBUTIONS

N.J-Y, S.R. conceptualization, supervision, funding acquisition, manuscript writing; E.Y, supervision, funding acquisition, manuscript writing; P.P.D, A.K., C.R, J.N.H, D.W., E.W.O, Q.T., Y.L.C., S.P.H., D.R., and K.K. experimentation, data analysis, generation of reagents M.I.C, C.K. single cell sequencing, C.P.N, D.L. scanning electron microscopy, all authors: editing of the manuscript.

## DECLARATION OF INTERESTS

The authors declare no competing interests.

## Notes

### Competing Interest Statement

The authors have declared no competing interest.

