## Supplementary figures and material and method for "Diversity and Function of Motile Ciliated Cell Types within Ependymal Lineages of the Zebrafish Brain"

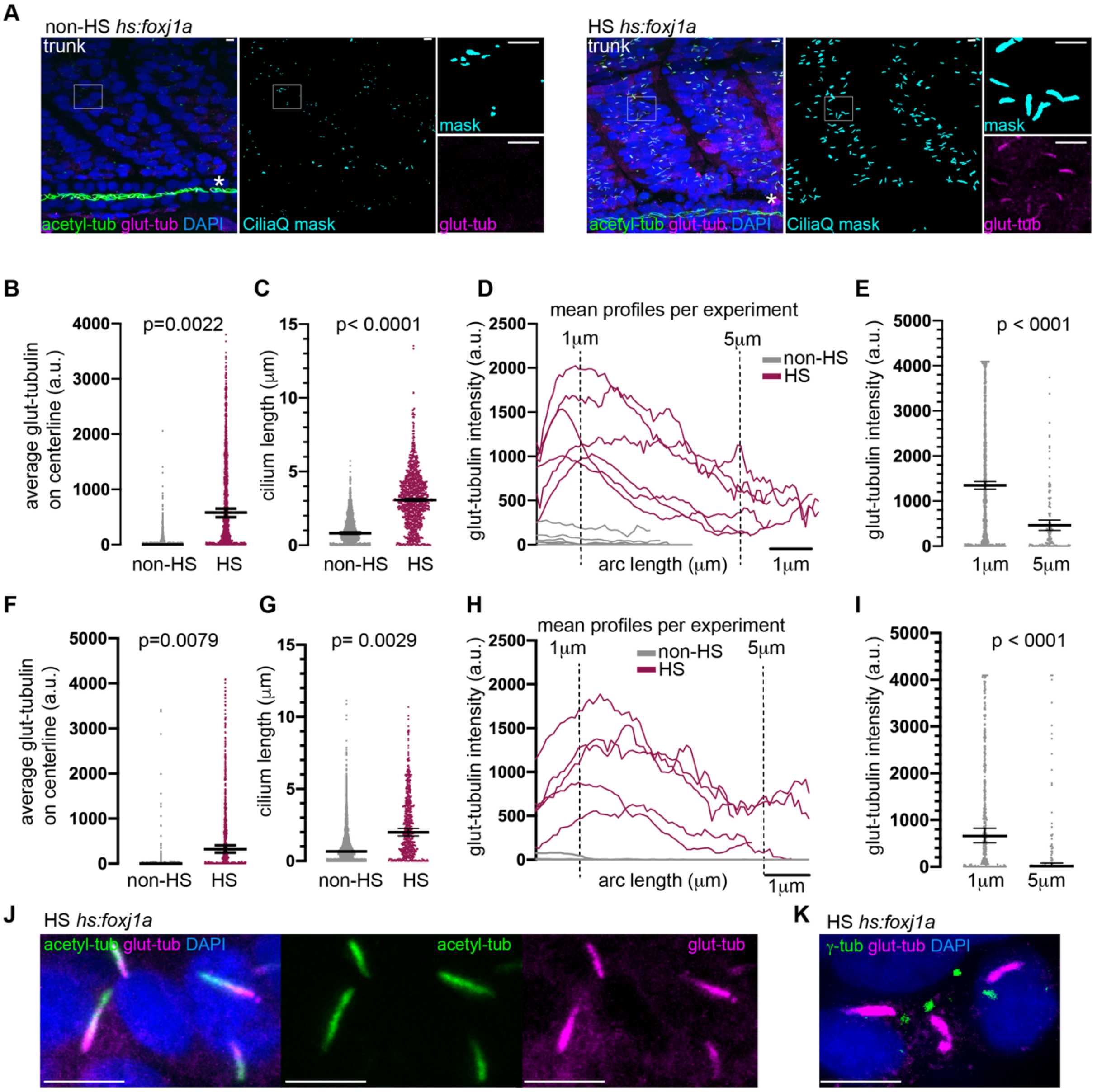


***Figure S1. Glutamylated tubulin is enriched in cilia of foxj1 expressing cells.***

***(A)*** *Confocal image of the trunk of a control (non-HS, left) and heat-shocked (HS, right) Tg(hs:foxj1a) transgenic embryo at 24 hpf immunostained with acetyl-tubulin (green), glutamylated tubulin (magenta) and DAPI. The acetyl tubulin signal was used for masking cilia using the CiliaQ software. Insets reveal longer and glutamylated-tubulin positive cilia in HS as compared to non-HS control. Representative example of n=6. The pronephros, indicated with *, was excluded from the analysis.* ***(B, F)*** *Overexpression of foxj1a by HS increases tubulin glutamylation in cilia in the trunk (B) and eye region (F). Each dot represents an individual cilium collected from all images, bars: median +- 95% confidence interval of median of all cilia. P-value by two-sided Mann-Whitney test on median of cilia from individual animals.* ***(C, G)*** *Overexpression of foxj1a in HS animals increased cilium length in trunk (C) and eye region (G). Each dot represents an individual cilium, bars: median +- 95% confidence interval of median of all cilia. p -value by unpaired, two-sided t-test with Welch correction on median of cilia from individual animals.* ***(D-E, H-I)*** *The intensity of glutamylated tubulin staining is not uniform, but enriched at one end of the cilium in the trunk (D-E) and eye region (H-I).* ***(D, H)*** *Each line represents the mean levels of glutamylated tubulin as a factor of position along the ciliary axoneme per embryo.* ***(E, I)*** *Each dot represents the levels of glutamylated tubulin at 1 μm or 5 μm along the ciliary axoneme for all individual cilium, bars: median +- 95% confidence interval of median. p with two-tailed Mann-Whitney test.* ***(J)*** *Glutamylated tubulin (magenta) is enriched at one end of the cilium as compared to* *acetyl tubulin (green), as shown in a representative confocal image of a 24 hpf heat-shocked Tg(hs:fox1a) embryo.* ***(K)*** *Glutamylated tubulin (magenta) is enriched at the base of the cilium as shown upon co-staining with the basal body marker gamma-tubulin (green) in a representative confocal image of a 24 hpf heat-shocked Tg(hs:fox1a) embryo. Scale bars are 5 µm.*

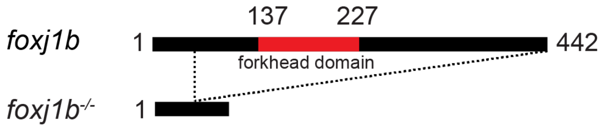


**insertion in mutant**
*deletion in mutant*

ATGCCGGTGTTAATGAGTCCTGAAATAGCGAATAAATTCAAAGAGAAATGGCTGATGCTTCATCCGGAGGATCAGGA
TAATGTCAGCGGCTCTGTGCACTTTGACGACAGTCTCACTAGTCTGCACTGGCTGCAGAACTTCTCCATCCTCAGT
GCCAACCCCGAGAGGACTCCCAGCTCCGGCTGCCACCCGCAACACCTCTTTTACTACAAAAACCAGCTGGGGGGT
ACCGACTCGCCCTCCAGTCCACCTGCCGGAGATACCGCCGCCACAGGGATGCCACAGACACCTGGGAATCCCAC
AACGTCCTGCAGCAGTTTGGCCAATCCATACGCGCTTCAGCAAGCCGGACACTACATA**TTAA***ACGGGGCAAACAAA*
*CCCGGCTGAGGAGATAGACTATAAAACAAACCGCCACGTGAAGCCACCCTATTCATATGCCACTCTGATTTGCATGG*
*CAATGCAAGCCAGCAACAAGACCAAAATCACCCTGTCAGCCATATACAGCTGGATCACCGAGAACTTCTGCTACTAC*
*AGATACGCAGAGCCGAGCTGGCAGAACTCAATCCGCCACAACTTGTCCCTGAACAAGTGCTTCATGAAAGTGCCC*
*AGGCAGAAAGACGAGCCAGGAAAAGGAGGCTTTTGGCAAATCGACCCTCAGTACGCTGACATGTTTGTGAATGGC*
*GTTTTCAAGAGACGGCGAATGCCTGCCACAAACTTCAATACCCAGAGACAAAGCAAAATGCTTTCCTCCCCGAGCT*
*CCTCGTACACCTCCCAGTGCAACCAACAAATGGGCATGGGCCACTTCCAAGGAAACAAACGCAAGCAAGTCTTCC*
*CCAAACGCGGGAACAAGCTAGCCCGTATCTCCAAGAGCCCTCTGTTAACCAGTGACATCAAAACCTCAGACGTCCT*
*GAGAGGAGACTTTGACCTGGCGTCGGTTTTCGATGACGTCCTAAGTGGGAATGACAGCACGTTTGAGGATTTGGAT*
*ATCAACACAGCTCTAAGCTCGCTAGGATGCGAAATGGAGCCGTCTTCCCAGATCCACAACCAGTCTGGTTATAGCA*
*ACGAGGATGAGCAGGCTTGTGCTTACCTGGAGGCCAACGGCATTATTGGATGCAACATGGAGGACTTCCACCACC*
*AGCAGCACCACCAGCAAGCACAGGTCCACCCG*CAATATTACGAGGGGATGTTCTCGGATCAGCAGAATCAACAACA
TCCTTGGGAGATTAAAGAGGAAGCCCAGCCAGTCCCTCTTTCGTTGGATCACGGCTATGCGTTTTGTGAGGGATTC
TTCTCAGAGATGCAACTTTGGGAAAGAGGAGAGTCATATATGTGA

***Figure S2. Schematic representation of the foxj1b$^{sq5719}$ mutant allele.***
A large part of the coding sequence of the *foxj1b* gene is lacking in the mutant allele, which includes the forkhead domain. The allele contains a small insertion (indicated in bold green) and a large deletion (indicated in italic red) of the gene.

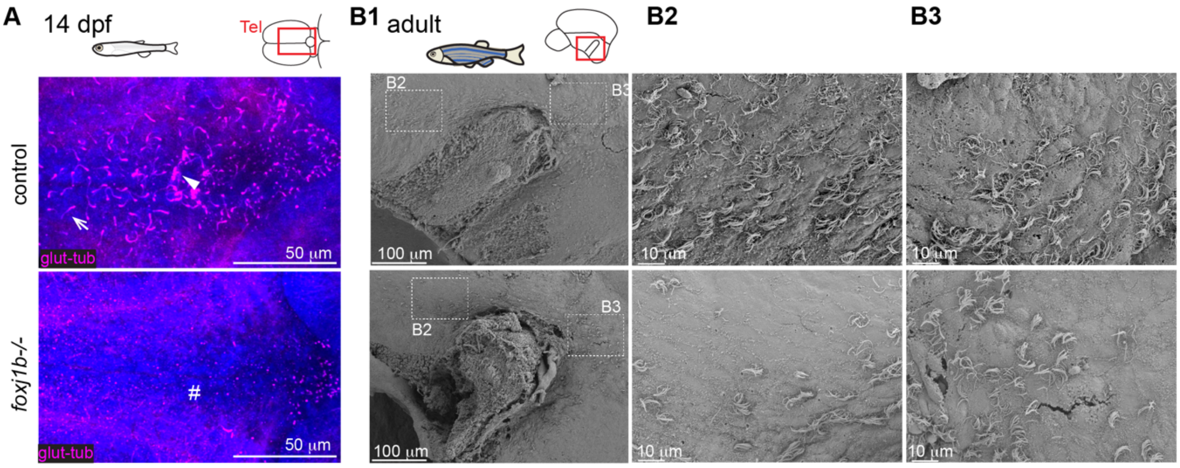


***Figure S3. Ciliary defects in foxj1b mutant in larval telencephalon and adult telencephalic midline.*** ***(A)*** *Confocal images at 2 weeks of age reveal that cilia are absent in the dorsal telencephalon of foxj1b mutant. Note the MCCs in the control (arrowhead). Cilia loss is indicated by #. (n=4).* ***(B1-B3)*** *Ciliated cells remain in the midline of the telencephalon of foxj1b mutants as shown by SEM analysis (n=3).* ***(B2-B3)*** *Magnified insets as shown in B1.*

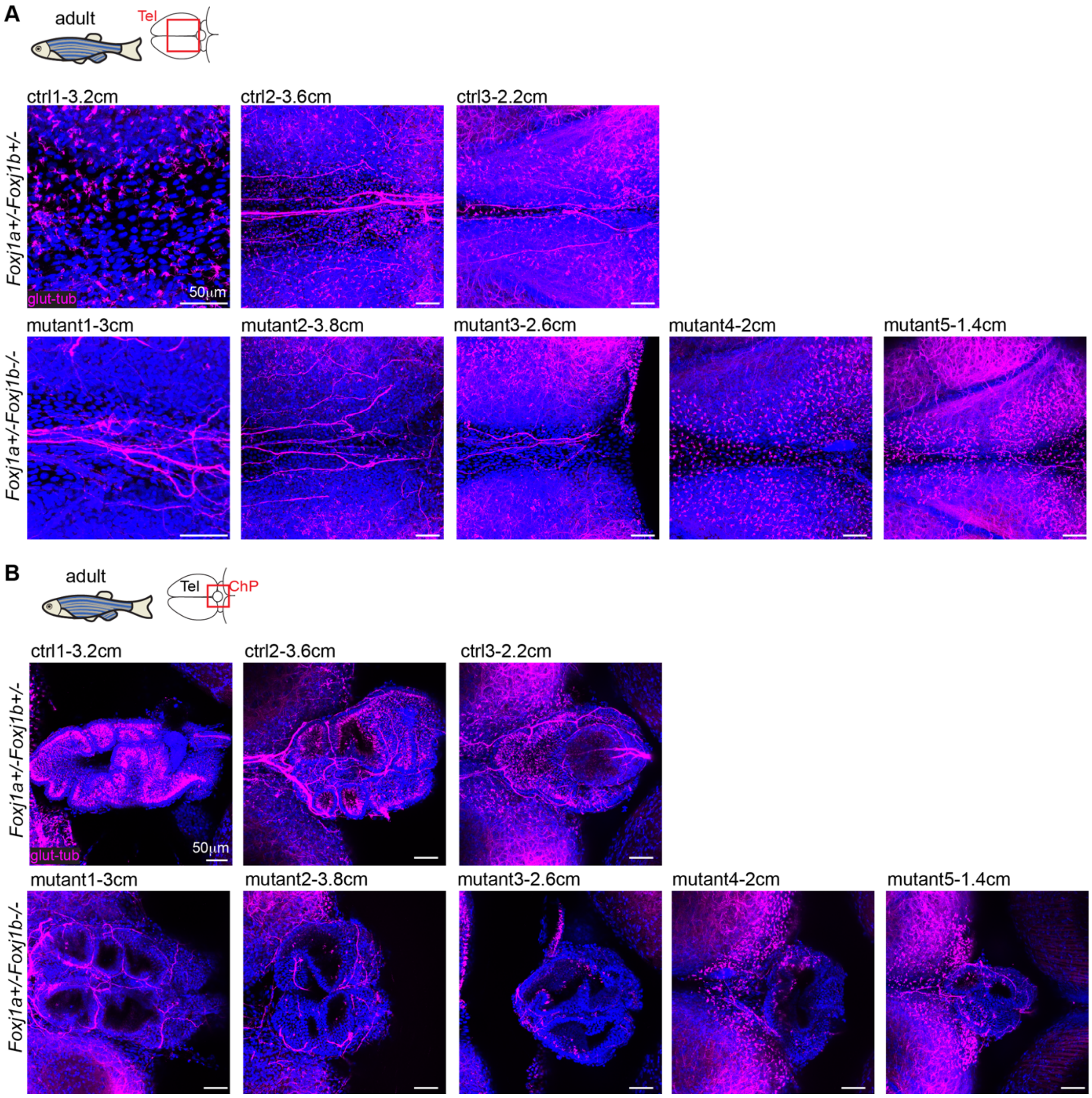


***Figure S4. Ciliary defects in the brain of foxj1a+/-;foxj1b-/- mutants are variable and more prominent in larger and older animals.***

*Confocal staining of adult foxj1a+/-;foxj1b+/- and foxj1a+/-;foxj1b-/- immunostained for glutamylated tubulin (magenta) and DAPI (blue) show a partial loss of MCC and total loss of single cilia in the TC **(A)** and the ChP (**B**) in foxj1a+/-;foxj1b-/- as compared to foxj1a+/-;foxj1b+/- controls. The effects are variable and more pronounced in larger/older animals. Body size (in cm) is indicated. Ctrl2 and mutant2 are included in Figure 6.*

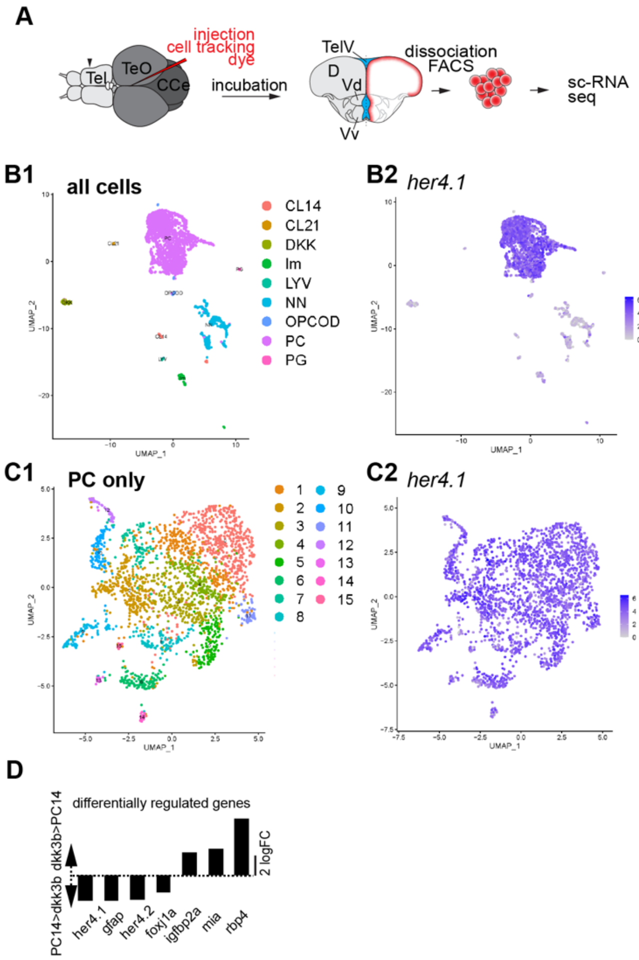


***Figure S5. Diversity of motile ciliated cells in the adult telencephalon revealed by single cell RNA seq analysis.***

*__(A)__ Single cell transcriptomic analysis was performed on cells lining the telencephalic ventricle upon injection with a cell tracking dye. A total of 3158 cells were analysed. __(B1-B4)__ Unbiased clustering analysis of 3158 sequenced cells revealed the presence of 9 cell clusters __(B1)__ expressing different cell markers, such as her4.1 expressing progenitor cells (PC, __B2__), SV2a expressing neurons (NN, __B3__), dkk3b expressing cells (DKK, __B4__). Im: immune cells, LYV: lyve1-expressing cells, OPCOD: oligodendrocyte cells/progenitors, PG: pineal gland. Individual dots represented on these tSNE plots correspond to a single cell. Blue intensities indicate expression levels. Grey dots denote cells that do not express the specified gene. __(C1-C4)__ The progenitor cell cluster (PC) was re-clustered into 14 different clusters expressing her4.1 broadly __(C2)__. A small percentage of cells expressed the proliferative marker pcna __(C3)__ in contrast to the quiescent marker s100b __(C4)__. __(D)__ Differentially regulated genes between the two ependymal clusters, PC14 and DKK clusters.*

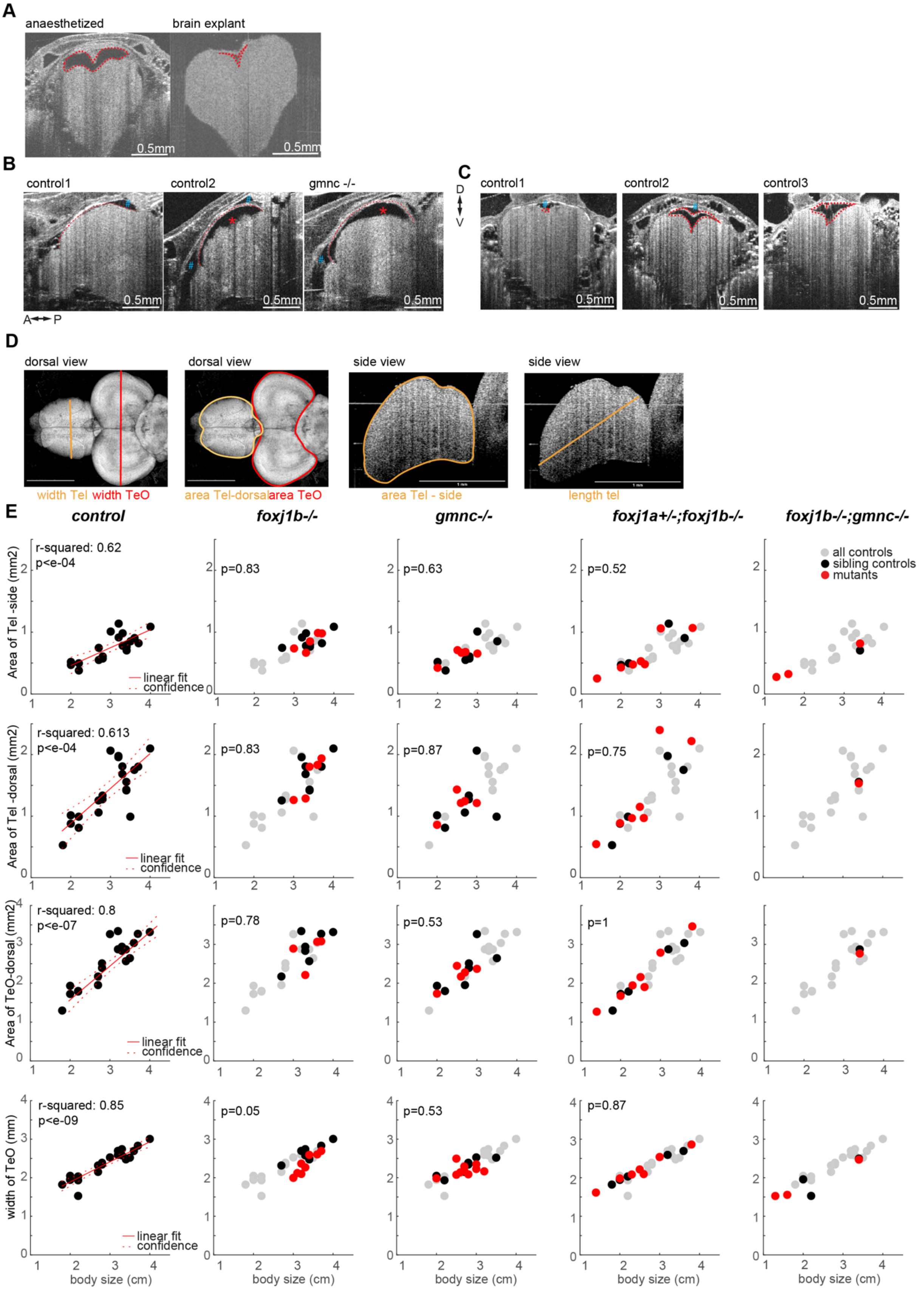


***Figure S6. Imaging of brain ventricles and brain explants by OCT.***
*(**A**) The telencephalic ventricle (indicated in red) collapses upon brain dissection. (**B**) Liquid is present both within the ventricle (indicated with a red *) surrounded by the TC (indicated with a red*

*dashed line), and within the skull (indicated with a blue #) outside the TC. Expansion of the telencephalic ventricle results in a reduction in volume of extra-ventricular liquid. (**C**) Ventricles of control animals are of different sizes. Three examples are shown.* ***(D)*** *Scheme indicating the different parameters quantified.* ***(E)*** *Quantifications of brain sizes represented as a function of body length. Left: all controls pooled. The linear relationship between body size and the various parameters was calculated using a linear model fit and reported with r-squared and p-values. The linear fit is indicated in red with the confidence interval. Others: all controls in grey, sibling controls in black, mutants in red. P-value (ranksum between sibling controls and mutants) are indicated in the graph. No p values are indicated for foxj1b;gmnc double mutants due to small sample size.*
*D: dorsal, V: ventral, A: anterior, P: posterior, TeO: optic tectum*

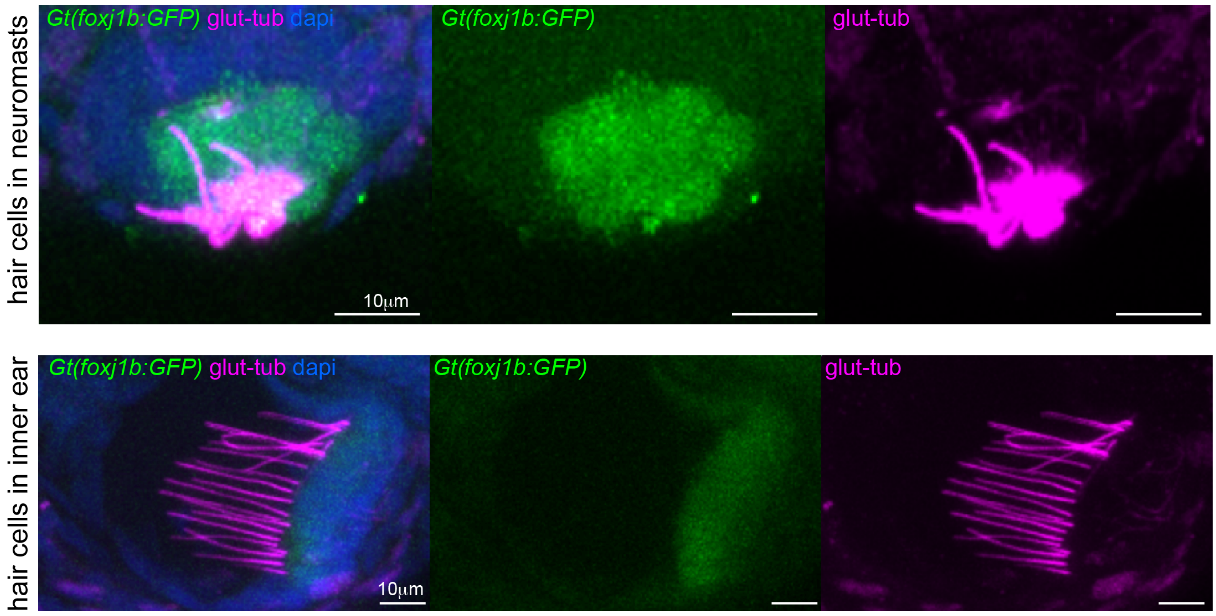


***Figure S7. Glutamylated tubulin is enriched in cilia of hair cells in neuromasts and inner ear.*** *Confocal images of Gt(foxj1b:GFP)$^{tsu10Gt}$ transgenic line immunostained for glutamylated tubulin (magenta) at 4 days (dapi in blue). Hair cells of neuromasts (top) and of inner ear (bottom) harbor glutamylated positive cilia.*

**Material and Methods**

| Reagent or Resource | Source | Identifier |
|---|---|---|
| **Antibodies** | | |
| 1. Mouse monoclonal glutamylated tubulin (GT335) | Adipogen | Cat#AG-20B-0020-C100; RRID: AB_2490210 |
| 2. Anti-gfp | Thermofisher | Cat # A-21311 |
| 3. Alexa fluor plus Goat anti mouse 488 | Thermofisher | Cat # A32723 |
| 4. Alexa fluor plus Goat anti rabbit 555 | Thermofisher | Cat # A32732 |
| 5. Beta catenin | cell signaling technologies | Cat#9562 |
| 6. Alexa fluor plus Goat anti mouse 555 | Thermofisher | Cat # A32727 |
| 7. Dapi | Thermofisher | Cat#D1306 |
| **Chemicals** | | |
| 1. x2 power up SYBR master mix | Thermofisher | Cat#15366158 |
| 2. Microamp optical 96 well reaction plate (Applied biosystems) | Thermofisher | Cat# N8010560 |
| 3. 70 kDa rhodamine B isothiocyanate-dextran | Sigma-Aldrich | Cat#R9379 |
| 4. SPHERO Fluorescent Yellow Particles 1% w/v, F = 1 µm | Spherotech | Cat#FP-1552-2 |
| 5. Phosphate buffered saline(10209252) | Thermofisher | Cat#BR0014G |
| 6. Triton X-100 | Merck | Cat# 1086031000 |
| 7. Bovine Serum Albumin (BSA) | PanReac AppliedChem | Cat#A1391 |
| 8. Dimethyl sulfoxide (DMSO) | Sigma | Cat#D8418 |
| 9. Glycerol | VWR | Cat#24387.292 |
| 10. Trizol | Thermofisher | Cat#15596026 |
| 11. Acetone | VWR | Cat#20066.296 |
| 12. Formaldehyde solution (PFA) | Sigma | Cat#F8775-25ml |
| 13. MS-222 | Sigma | Cat#E10621-60G |
| 14. Proteinase K from tritirachium album | Sigma | Cat#P2308-25MG |
| 15. Tris | Sigma | Cat#252859-100G |
| 16. RevertAid First Strand cDNA Synthesis Kit | Thermofisher | Cat#K1622 (100 reactions) |
| 17. CellTracker™ Red CMTPX Dye | Thermofisher | Cat# C34552 |
| **Oligonucleotides** | IDT | |
| Gmnc genotyping crispr deletion Forward TTGTGATTGTCTCATGCTGTTG | IDT | |

| | | |
|---|---|---|
| Gmnc genotyping crispr deletion Reverse AAAAATTCCAGTTTGTCAAGGC | IDT | |
| Foxj1b genotyping crispr deletion Reverse CTCCATCCTCAGTGCCAACC | IDT | |
| Foxj1b genotyping crispr deletion Hom CGGCTCTGCGTATCTGTAGT | IDT | |
| Foxj1b genotyping crispr deletion Forward TCTTCAGACCAGCAAAGACAGT | IDT | |
| Foxj1a Reverse genotyping CGCTATCGAGGAAGGACAGGATTT | IDT | |
| Foxj1a Forward genotyping GCTGGTCAGGCTGTCGTCTAAA | IDT | |
| Foxj1b Reverse GGGCACTTTCATGAAGCACT | IDT | |
| Foxj1b Forward ACAGCTGGATCACCGAGAAC | IDT | |
| Foxj1a Forward TTTAGACGACAGCCTGACCA | IDT | |
| Foxj1a Reverse GGTCTCCTGCTAACGGTGAA | IDT | |
| **Software and Algorithms** | | |
| ImageJ/Fiji | | [96] |
| CiliaQ | Wachten lab, University of Bonn | [97] |
| **Zebrafish lines** | | |
| *T2BGSZ10 Gt(Foxj1b:GFP)* | Meng lab, Tsinghua University | [98], ZFIN: ZDB-ALT-110301-1 |
| Gmnc mutant (*gmnc*$^{sq34}$) | Roy lab, A*STAR | [59] ZFIN: ZDB-ALT-160901-7 |
| Foxj1b mutant (*foxj1b*$^{sq5719}$) | Roy lab, A*STAR | This study |
| Foxj1a mutant (*foxj1a*$^{nw3}$) | Yaksi lab, NTNU | [14] ZFIN: ZDB-ALT-190620-14 |
| *Tg(foxj1a:gfp)BAC*$^{vcc41}$ | Kikuchi lab, NCVC | This study |
| *Tg(Hs:foxj1a)*$^{sq5713Tg}$ | Roy lab, A*STAR | [39], ZFIN: ZDB-ALT-141111-1 |
| **Other** | | |
| Pressure injector | Eppendorf | Femtojet 4i |
| Confocal microscope | Zeiss | Examiner Z1; Olympus Fluoview |
| Optical Coherence Tomography | Thorlabs | Telesto 1300nm SD-OCT and LK-4 objective |
| Step One Real Time PCR system | Thermofisher | Catlog#4376357 |

**Zebrafish maintenance and strains:**

The animal facilities and maintenance of the zebrafish, *Danio rerio*, were approved by the NFSA (Norwegian Food Safety Authority) and the Singapore National Advisory Committee on Laboratory Animal Research. All the procedures were performed on zebrafish larvae of different developmental stages post fertilization in accordance with the European Communities Council Directive, the Norwegian Food Safety Authorities and the Singapore National Advisory Committee on Laboratory Animal Research. Embryonic, larval and adult zebrafish were reared according to standard procedures of husbandry at 28.5 °C, unless mentioned otherwise. For our experiments, the following fish lines were used: *T2BGSZ10 Gt(foxj1b:GFP)* [98], *gmnc* mutant [59], *foxj1a* mutant [14], *Tg(hs:foxj1a)* [39], *foxj1b* mutant and *Tg(foxj1a:gfp)* (see below).

**CRISPR/Cas9 mediated mutation of *foxj1b***

For generating a mutation in the *foxj1b* gene, we used two gRNAs targeting regions within exon 2 and exon 3. The gRNA sequences are as follows: Exon 2 5'-AAGCCGGACACTACATAACG-3' and exon 3 5'-GGTCCACCCGCAATATTACG-3'. This resulted in a large deletion of 804 nucleotides, as well as addition of 4 nucleotides at the 5' gRNA target site in exon 2. As a consequence, the mutant Foxj1b protein is expected to be highly truncated and consist of only 140 amino acids (bereft of the forkhead as well as the transactivation domains), of which 120 retains identity with wild-type Foxj1b, and is likely to be completely non-functional.

**Generation of *TgBAC(foxj1a:gfp)$^{vcc41}$* transgenic strain**

The *TgBAC(foxj1a:EGFP)$^{vcc41}$* construct was generated by inserting the EGFP expression cassette into the CH211-151D17, after the *foxj1a* translational start codon, using Red/ET recombineering (GeneBridges, Heidelberg, Germany). The final construct was purified using the BACMAX DNA Purification kit (Epicentre, Madison, WI, USA), linearized with *Sfi*I, and injected into the single cell-stage zebrafish eggs to establish stable transgenics.

**Genotyping**

For genotyping, the samples were subjected to gDNA isolation using 100 µl PCR lysis buffer (containing 10 mM tris pH7.5, 50 mM EDTA, 0.2% tritonX-100 and 0.1mg/ml Proteinase K) overnight at 50 °C. To stop the reaction the samples were heated to 95°C for 10 minutes. The samples were then centrifuged at 13000 rpm for 2 minutes. The supernatant containing gDNA was used for qpcr. For performing qpcr, 5 µl SYBR green pcr master mix (Thermofisher) was mixed with 0.5 µl each of forward and reverse primer and 4 µl water to make a 10 µl reaction mixture. This reaction was added to a 96well qpcr plate (Thermofisher) and 2 µl of extracted gDNA were mixed with this reaction. The samples were then analyzed based on their melting curves as wt, het or homozygous.

**RNA extraction, cDNA preparation and qPCR**

To prepare cDNA for qpcr, RNA was individually isolated from dissected tissues pooled from 6 animals. The tissues analyzed included the tela choroida, telencephalon and rest of the brain. The samples were first homogenized using 500 µl Trizol (Thermofisher, cat number 15596026). The samples were homogenized using a 27G needle to make a uniform solution. After homogenizing the solution, another 500 µl of trizol was added and incubated for a period of 5 minutes. Post incubation 200 µl of chloroform was added and the tube was mixed well for 15 seconds and incubated for 2 minutes at RT. The tubes were then centrifuged at 4°C for 15 minutes. After centrifugation, the mixture separated into lower red phenol-chloroform phase, an interphase and an upper colorless aqueous phase. This upper aqueous phase was carefully transferred into a fresh tube. 500 µl Isopropanol and 0.5 µl RNAse free glycogen (Thermofisher) were added to the mixture and mixed thoroughly. The mixture was then incubated at 4°C for 10 minutes. After incubation the samples were centrifuged at 4°C for 10 minutes. The supernatant was discarded, and the pellet was washed with 1ml of cold 70% ethanol. The samples were centrifuged at 4°C for 10 minutes. The supernatant was discarded, and the pellet was air dried at RT for 10 minutes. After air drying the pellet was resuspended in 15-20 µl RNAse free water and measured for its concentration and purity using a Nanodrop. For making cDNA a ReverseAid first strand cDNA synthesis kit (Thermofisher) was used after DNase treatment. First, 1 µg of total RNA was treated with 1 µl of DNase at 37°C for 30 minutes. After incubation, 1 µL of EDTA (50mM) was added to the mixture and incubated for 10 minutes at 65°C to inactivate the DNase enzyme. After this step, 1 µl of Oligo $(dT)_{18}$ primer was added to the reaction and incubated for 5 minutes at 65°C and then 5 minutes at 4°C. To complete the synthesis step, 4 µl of 5X reaction buffer, 1 µl of Ribolock RNase Inhibitor (20

U/µl), 2 µl of 10 mM dNTP Mix and 1 µl of RNase-free water or RevertAid RT (200 U/µl) were added and the mixture was gently centrifuged. This was then incubated at 42°C for 60 minutes and then 70°C for 5 minutes. The final cDNA mixture was diluted using 100 µL RNase free water.
For performing qpcr a total of 10 µl reaction per well was prepared (master mix solution of 5 µl of x2 power up SYBR was mixed with 0.5 µl each of forward and reverse primer (10µM) and 4 µl of milliQ water). This solution was added to individual wells and 2 µl of cDNA sample was added to each well in duplicates. The plates were run using the following Run method. The holding stage was set at 95°C for 10 mins, the cycling stage was set at 95°C for 15 mins followed by 60°C for 1 minute (40 cycles in total). The melt stage was set using 3 steps of 95°C for 15 mins, 60°C for 1 minute followed by another step of 95°C for 15 mins using a 0.1°C increment in step 3.

**Ectopic expression of *foxj1a* by heat shock**

*Tg(Hs:foxj1a)* embryos at ~18 hpf were heat-shocked at 37°C in water bath for one hour. The heat-shocked embryos were further incubated at 28°C until 24 hpf, before proceeded for fixation and antibody staining. Controls included both non-heat shocked transgenic as well as heat shocked non-transgenic wild-type.

**Antibody staining and confocal imaging**

**Immunostaining of the brain**

Larvae or juvenile fishes were first euthanized and then fixed in a solution containing 4 % paraformaldehyde solution (PFA), 1 % DMSO and 0.3 % TritonX-100 in PBS (0.3 % PBSTx) for at least 2 h at room temperature or 4 degrees overnight. Stainings were performed on cut heads to improve the penetration of the antibodies. The larvae were washed with 0.3 % PBSTx after fixing to remove any traces of the fixing solution. For permeabilization, samples were incubated for 10 min (larvae) or 1h (juvenile/adult) at -20 °C (larvae) with acetone. Subsequently, samples were washed with 0.3 % PBSTx (3x10 min) and blocked in 0.1 % BSA/0.3 % PBSTx for 2 h (larvae) or 1% BSA in 0.3% PBSTx for 4 hours (adult) at room temperature. Samples were incubated with glutamylated tubulin (GT335, 1:400, Adipogen) for staining cilia and beta-catenin antibody (1:400, 9562-cell signalling antibodies) overnight at 4 °C. On the second day, samples were washed (0.3 % PBSTx, 3x1 h) and subsequently incubated with the secondary antibody (Alexa-labelled GAM488 plus, and GAM555 plus Thermo Scientific, 1:1,000) and 0.1 % DAPI overnight at 4 °C. For gfp staining ( Anti-gfp tagged polyclonal antibody alexa fluor 488,Thermo Scientific) was used to enhance the GFP signal if needed. On the third day after incubation with the secondary antibody the larvae were washed (0.3 % PBSTx, 3x1 h) and transferred to a series of increasing glycerol (made in PBS) concentrations (25 %, 50 % and 75 %). After staining the larvae were stored in 75 % glycerol at 4 °C and imaged using a Zeiss Examiner Z1 confocal microscope with a 20x plan NA 0.8 objective.

**Immunostaining on *foxj1a* over expressed embryos**

The following primary antibodies were used: rabbit anti-acetylated tubulin (1:500, Cell Signaling), mouse anti-glutamylated tubulin (1:500, Adipogen). Primary antibodies were diluted in PBDT (1% (w/v) BSA, 1% DMSO, 0.5% Triton X-100, PBS base), and incubated with the embryos at 4 °C for overnight. After extensive washes with PBDT, the embryos were then incubated with Alexa Fluor-conjugated secondary antibodies (Invitrogen, 1:500) at 4 °C for overnight. DAPI staining were subsequently performed at room temperature for 30 min, and the stained embryos were thoroughly washed with PBDT, before mounted in 70% glycerol. Confocal imaging was performed with Olympus Fluoview Upright Confocal Microscope, with a 100x plan NA 1.45 objective.

**CiliaQ analysis**

Confocal images were analyzed using the CiliaQ workflow [97, 99]. First, images were pre-processed and segmented with CiliaQ Preparator (Settings: Gaussian Blur with sigma 1.0, Hysteresis threshold with low threshold delivered by algorithm Huang and high threshold delivered by algorithm Triangle, Threshold determined in a maximum-intensity-projection). Second, images were edited with CiliaQ Editor by a trained observer: incompletely segmented cilia were connected manually, multiple adjacent cilia connected to one object were separated, long motile cilia at the pronephros were removed from the analysis. Third, images were analyzed with CiliaQ to quantify the cilia in the images (settings: minimum cilium size 25 voxel, cilia touching X Y or Z borders excluded, Gauss XY sigma 2.0 voxel, Gauss Z sigma 0.0 voxel). Because pixel size differed between images (either 0.124 µm or 0.079 µm, stack interval in all images: 0.4 µm), profiles from images with higher pixel size were scaled to the lower pixel size by linear interpolation (using the R function approxfun()). Next, Ciliary glutamylated-Tubulin intensity profiles were aligned with a custom-written algorithm in R. The integrated intensity of the first third and the last third of the CiliaQ-derived intensity profiles was compared. If the integrated intensity of the last third was higher than the integrated intensity of the first third, the profile was reversed. Graphical representations and statistical analysis were done using GraphPad. The type of statistical test performed depended on the results from Normality and Lognormality tests (Shapiro-Wilk, Kolmogorov-Smirnov) were passed is indicated in the respective figure legend

**Brain ventricle injections and imaging**

Prior to injections, all larvae/juveniles were anaesthetized in 0.01% MS-222 in AFW. Injections were done on larvae/juvenile embedded in 2% low-melting point agarose in AFW and 0.01% MS-222 in AFW. For animals older than 2 months, animals were first euthanized by hypothermia. The brain was then dissected in artificial cerebrospinal fluid (aCSF) and mounted on sylgard using a metal pin [100, 101]. The ACSF was composed of the following chemicals diluted in reverse osmosis–purified water: 131 mM NaCl, 2 mM KCl, 1.25 mM $KH_2PO_4$, 2 mM $MgSO_47H_2O$, 10 mM glucose, 2.5 mM $CaCl_2$, and 20 mM $NaHCO_3$. The injection mixture contained 70 kDa rhodamine B isothiocyanate-dextran (RITC-dextran; Sigma-Aldrich, R9379) dissolved in aCSF at a final concentration of 10 mg/ml. The needles used for the injections were pulled with a Sutter Instrument Co. Model P-2000, from thin-walled glass capillaries (1.00 mm; VWR®), using the following settings: heat = 785, filament = 4, velocity = 40, delay = 220, pull = 70. The needle tip was cut open with forceps. A pressure injector (Eppendorf Femtojet 4i) was used to inject between 1 (larvae) -15(adult) nl of solution (depending on the developmental stage) in the telencephalic ventricle. The pressure and time used for the injection were calibrated for each needle using a 0.01 mm calibration slide for microscopy. After injection, the larvae, juvenile or brain explant were immediately transferred to the confocal microscope (Zeiss Examiner Z1), and imaged with a 20x water-immersion objective (Zeiss, NA 1.0, Plan-Apochromat) at room temperature. In order to avoid bodily movement during the recordings, larvae/juvenile were maintained in AFW containing 0.01 % MS-222. The laser power was corrected according to the depth of imaging. Multiple images were acquired per sample and stitch using the 3D stitching plugin in Fiji/ImageJ. Prior to 3D reconstruction, the 3D stacks were gaussian blurred and reduced in size. 3D reconstructions were done using the 3D viewer plugin in Fiji/ImageJ (threshold levels were adjusted for each sample), exported as stl file, and projected in 2D in Photoshop.

**Image processing**

To generate whole brain images shown in figure 1 and 2, confocal stacks were stitched in Fiji/ImageJ using the deprecated 3D stitching plugin [102]. For Figure 1A2, adjacent maximum projected stacks were aligned manually and blended using the auto-blend function in Photoshop. To avoid a pixelated appearance of zoom-in inset, the pixel resolution of the inset was increased in Fiji/ImageJ by doubling the pixel number. The colors for Figures 2D4-D5 were modified in Photoshop. Figures were assembled using Illustrator.

**Scanning electron microscopy**

For *gmnc* and *foxj1b* mutants, 6 sets of WT and mutant zebrafish brain samples (dissected midline along the sagittal plane) were fixed by immersion in 4% Formaldehyde and 2% Glutaraldehyde in 0.1M cacodylate buffer, post-fixed with 1% osmium tetroxide in distilled water and dehydrated in ethanol series up to 100% Ethanol. Dehydrated samples were dried using critical point drying (Leica EM CPD030), mounted onto aluminum stubs with double sided carbon tape and sputter coated with 4 nm layer of platinum (Leica EM SCD050). SEM analysis was performed using JSM 6701F SEM (JEOL) operating at 5kV.

**Recording of ciliary beating**

To measure ciliary beating in adult brain, we performed light transmission microscopy on a brain explant perfused with aCSF bubbled with carbogen (95% $O_2$/5% $CO_2$). Recordings were obtained using a Manta camera and a custom designed software as previously described [8]. Recordings were obtained on an upright Olympus microscope with a 40x water immersion objective at circa 100Hz frame rate for 30 s. The resolution with the 40x objective was 1pixel=0.314µm. Multiple recordings were performed per brain explant along the dorsal telencephalon or along the telencephalic midline. Next, we aligned all light-transmission brain ventricle recordings using a custom-written alignment algorithm[8], adapted from Bergen et al [103], which corrects for occasional x-y drifts. We assumed that beating cilia are manifested in the acquired recordings as periodic changes in pixel intensity. To extract those periodic changes, we performed the MATLAB fast Fourier Transform algorithm for every pixel in the recording. From the resulting power spectra, we determined the primary frequency as the frequency of the highest peak between a lower frequency cut off (see table) and half the frequency of acquisition. Those primary frequencies were then rendered in the original pixel locations to form a frequency map. Beating cilia do not cover the entire recording and should span a minimum number of pixels. To segment signal and noise regions, we used a custom standard deviation (SD) thresholding-algorithm. In brief, we moved a 3x3 kernel across the entire frequency map and any pixel belonging to a 3x3 kernel whose SD was below 2 (see table) was considered as signal, whereas pixels belonging only to kernels with an SD above 2 were considered as noise. To remove small signal islands, we listed all connected pixels in the frequency map using the *bwconncomp* Matlab function, and removed those islands that were smaller than 500 pixels (see table). Altogether, with SD-thresholding algorithm and by removing small signal islands, we were able to restrict the analysis only to those regions with beating cilia.
Since we recorded from opaque explanted tissue, we encountered several additional sources of noise that would hinder frequency detection: not all movement could be resolved by alignment; wandering tissue particles occasionally interfered with the ciliary signal; and the opaque tissue limits the detection of a ciliary signal such that that its peak frequency is overshadowed by a power spectral low frequency shoulder, reminiscent of pink noise [104].
To ensure the origin of a signal, we interpreted the raw recordings. All ciliary signals should be constituted by visible ciliary beating. When a signal did not meet this requirement, we attempted to remove it by either pre-processing the recording with a high-pass filter – designed using *designfilt* and applied with *filtfilt* in Matlab (see table) - or by manually masking a spatially separated noisy signal using a custom-written script in Matlab. Occasionally, we detected signals from two distinct brain regions in one recording. Here, we segmented the recording as well. Recordings without a ciliary signal or with remaining noise were excluded from the analysis.
For a given recording, we collected the primary frequencies in a histogram and normalized its count with the number of signal pixels. We first averaged all histograms for a given brain region and fish, and then across fish to obtain one histogram per brain region: tela choroida, choroid plexus, and midline.

*Settings*

| | **Dura** | **Midline** | **Choroid Plexus** |
|---|---|---|---|
| **Lower frequency cut off** | 5.5 [Hz] | 10 [Hz] | 10 [Hz] |
| **Standard deviation** | 2 | 2 | 2 |
| **Spatial resolution** | 0.31445 [µm / pixel] | 0.31445 [µm / pixel] | 0.31445 [µm / pixel] |
| **Minimum number of pixels** | 500 | 500 | 500 |

*Table 1: Filter specifications*

| | **Dura** | **Midline** | **Choroid Plexus** |
|---|---|---|---|
| **Type** | highpassiir | highpassiir | highpassiir |
| **StopbandFrequency** | 2 | 4 | 4 |
| **PassbandFrequency** | 5 | 10 | 10 |
| **StopbandAttenuation** | 2 | 2 | 2 |
| **PassbandRipple** | 1 | 1 | 1 |
| **Sample Rate*** | ~100Hz | ~100Hz | ~100Hz |

| | | | |
|---|---|---|---|
| **DesignMethod** | butter | butter | butter |
| **MatchExactly** | passband | passband | passband |

* varies slightly per recording

To produce the Figure 3B1, C1, D1, transmission microscopy and ciliary beating heatmaps images were generated by Matlab in png format from individual recordings. The transmission microscopy images were first aligned manually in Photoshop using overlapping information from neighboring images. Then the ciliary beating frequency map were aligned to the transmission microscopy images and their background was removed using blending options in Photoshop.

**Recording of fluid flow**

To measure the directionality of fluid flow, we injected 1µm beads in the telencephalic ventricle of a brain explant as described above. The injection mixture contained fluorescent beads (SPHERO Fluorescent Yellow Particles 1% w/v, F = 1 mm) diluted in 1/6 in RITC-dextran containing aCSF. Fluid flow was measured using an epifluorescence microscope, a 10x water immersion objective and an ocular camera using an acquisition speed of 24-50 frames per second for at least 1 minute. The resolution with the 10x objective was 1pixel=1,13µm.
The particles were tracked using the trackmate plugin in Fiji/ImageJ [105] using the LoG detector for 8-10 pixel blobs and a threshold of 10-50 depending on the recording. Other parameters were initial search radius: 15,0, max search radius: 15,0, max frame gap: 2. The tracks were then imported in Matlab. Only tracks containing more than 20 data points and covering a distance longer than 10 pixels were included in the further analysis. The direction between two consecutive data point was then plotted using the surf function in Matlab.

**Single-cell sequencing, reads alignment and analyses**

2 female and 1 male 12 months old fish were injected with cell tracking dye (CMPTx, Cat# C34552; Invitrogen) [64]. 5-6 min post-injection, fish were euthanized and telencephalon dissected followed by dissociation and Fluorescent Associated Cell sorted [64, 107].
To prepare the cells for droplet-based sequencing, around 15000 cells were flow-sorted into a Bovine serum albumin (BSA) coated tubes containing 2 µl saline solution with 0.04 % BSA. Subsequently, the single cell suspension was carefully mixed with reverse transcription mix before loading the cells on the 10 X Genomics Chromium systems. During the encapsulation, the released polyadenylated RNA bound to the barcoded bead, which was encapsulated with the cell. Following the guidelines of the 10x Genomics user manual, the droplets were directly subjected to reverse transcription, the emulsion was broken and cDNA was purified using Dyna beads MyOne Silane beads. After the amplification of cDNA with 10 cycles, purification and quantification was performed. Half of the material of the cDNA was fragmented for five minutes and dA-tailed, followed by adapter ligation step and an indexing PCR of 10 cycles in order to generate libraries. After quantification, the libraries were sequenced on an Illumina NovaSeq machine using an S1 flowcell in PE mode (R1: 29 cycles; I1: 8 cycles; R2: 93 cycles).
The raw sequencing data was then processed with the 'count' command of the Cell Ranger software (v3.1.0) provided by 10X Genomics. The option of '--expect-cells' was set to 5,500 (all other options were used as per default). To build the reference for Cell Ranger, zebrafish genome (GRCz11) as well as gene annotation (Ensembl 95) were downloaded from Ensembl and the annotation was filtered with the 'mkgtf' command of Cell Ranger (options: '--attribute=gene_biotype:protein_coding --attribute=gene_biotype:lincRNA –attribute=gene_biotype:antisense'). Genome sequence and filtered annotation were then used as input to the 'mkref' command of Cell Ranger to build the appropriate CellRanger Reference.
In order to analyze the single-cell data, in total 3808 cells recovered after filtering by cell ranger. Then, outlier cells with 5-fold difference between total number of reads (nCount_RNA) and total number of features (nFeature_RNA) were removed from analysis. We further removed cells; with less than 500 genes, with more than 6% mitochondrial RNA genes. The remaining 3158 cells were used for downstream analysis with Seurat (3.2.0). We performed 3 iterative clustering; (i) clustering of all cells to identify the main cell types as done previously [64], (ii) the clustering of Progenitor Cells (PC) to identify subtype of PC cells and (iii) the clustering of Foxj1 positive (*foxj1a* and/or *foxj1b*) cells.

All cells were clustered by scaling to 10e4 and using top 2000 variable genes. nCount_RNA and nFeature_RNA were regressed out and dims = 50 and resolution=1 were used to identify clusters. The following markers were used to identify main cell clusters; Progenitor Cells (PC) (*fabp7a*), Neurons (NN) (*sv2a*), Immune Cells (Im) (*lcp1, wasb, pfn1*), Oligodendrocytes (OPC) (*olig1/2, aplnra/b*), Pineal Gland (PG) (*exorh*), DKK (*dkk3b*), LYV (*lyve1*) and unassigned clusters (14 and 21; named as CL14 and CL21, respectively).
The PC cells were further clustered as above parameters; except with dims = 30. Foxj1 cells were similarly clustered, except dims = 30 and resolution = 0.6. Additionally, PC cells and DKK cells were clustered together, as DKK cells contains *foxj1a/b* and *gmnc* genes.
We use monocle2 [71] to perform cell trajectory for all clustering of PC and foxj1 cells as described previously [64]. In brief, we use the scaled data and variable genes from Seurat and set SizeFactor 1.0 to prevent any further normalization.
The raw data for single-cell sequencing have been deposited on Gene Expression Omnibus and will be made publicly available upon acceptance of the manuscript.

**OCT imaging**

Adult zebrafish of various genotype were anaesthetized with 0.01% MS222 in bubbled and pH adjusted (using bicarbonate buffer) artificial fish water. Following loss of balance, anaesthetized animals were placed in a plastic container and immobilized using wet kim wipes. Animals were imaged using a Telesto OCT (Thorlabs, laser 1300nm) with the LK04 objective (resolution of 20micron). Following image acquisition, animals were euthanized and the brain explanted prior to be re-imaged in the OCT. Quantification of brain size and ventricle area was measured using the polygonal selection tool and the straight-line selection tool on resliced images in Fiji. Linear regressions were calculated in Matlab using fitlm function.
Brains of 3 additional *foxj1b;gmnc* double mutants were dissected and inspected using a stereomicroscope. None of these animals showed major brain malformations or size differences. These quantifications were not included in figure 7 because photos were made in fixed samples using a stereomicroscope.
